# Caveolae couple mechanical stress to integrin recycling and activation

**DOI:** 10.1101/2022.04.27.489710

**Authors:** Fidel-Nicolás Lolo, Dácil M. Pavón, Araceli Grande-García, Alberto Elósegui-Artola, Valeria Inés Segatori, Sara Sánchez-Perales, Xavier Trepat, Pere Roca-Cusachs, Miguel A. del Pozo

## Abstract

Cells are subjected to multiple mechanical inputs throughout their lives. Their ability to detect these environmental cues is called mechanosensing, a process in which integrins play an important role. During cellular mechanosensing, plasma membrane (PM) tension is adjusted to mechanical stress through the buffering action of caveolae; however, little is known about the role of caveolae in early integrin mechanosensing regulation. Here, we show that Cav1KO fibroblasts increase adhesion to FN-coated beads when pulled with magnetic tweezers, as compared to wild type fibroblasts. This phenotype is Rho-independent and mainly derived from increased active β1-integrin content on the surface of Cav1KO fibroblasts. FRAP analysis and endocytosis/recycling assays revealed that active β1-integrin is mostly endocytosed through the CLIC/GEEC pathway and is more rapidly recycled to the PM in Cav1KO fibroblasts, in a Rab4-dependent manner. Moreover, the threshold for PM tension-driven β1-integrin activation is lower in Cav1KO MEFs than in wild type MEFs, through a mechanism dependent on talin activity. Our findings suggest that caveolae couple mechanical stress to integrin cycling and activation, thereby regulating the early steps of the cellular mechanosensing response.

## Introduction

Cells constantly adjust their PM composition in response to changes in extracellular matrix (ECM) stiffness, which regulates many aspects of cell behavior^1^. How cells sense and react to ECM stiffness changes is critical to understanding both physiological and pathological processes^2^. The ECM is mechanically linked to the cytoskeleton through integrins, which provide a bridge between extracellular cues and downstream cellular events^3^. The mechanosensing function of integrins is well established, and much progress has been made in defining the molecular details of integrin action; for example, how α5β1 and αvβ3 integrins withstand and detect forces, respectively^4^, and how differences in integrin bond dynamics contribute to tissue rigidity sensing^5^. Integrin function requires an activation step commonly achieved by binding to activator molecules, such as talin and kindlins^6^. Integrins are also activated in response to changes in membrane tension, as recently reported by Ferraris *et al.*^7, 8^. However, it is unclear how these events influence mechanosensing.

Tissues experiencing wide variations in PM tension, such as endothelium, muscle, fibroblasts, and adipocytes, have a high membrane density of caveolae^9–11^. Caveolae are 60-80 nm PM invaginations and are key elements in the sensing and transduction of mechanical forces. Core caveolae protein components are caveolin-1 (Cav1) and cavin-1 (also called PTRF), and lack of either results in caveolae loss^12, 13^. Cav1 is intimately linked to integrins^11, 14, 15^ and to molecules such as Filamin A, which links integrins to the cytoskeleton^16^. To study how integrins and caveolae interact to regulate cell mechanosensing, we used magnetic tweezers (MT)^17^ to exert mechanical force on mouse embryonic fibroblasts (MEFs) upon their binding to magnetic beads coated with the integrin-binding protein fibronectin (FN). MEFs lacking Cav1 (Cav1KO) showed higher adhesion to FN-coated beads than Cav1WT MEFs, through a process dependent on increased β1-integrin surface availability in Cav1KO MEFs due to both rapid recycling of endocytosed β1-integrin to the PM, in a Rab4-dependent manner, and tension-mediated β1-integrin activation driven by Talin. These results support a role for caveolae in regulating integrin mechanosensing by coupling membrane tension to integrin recycling and activation.

## Results

### Genetic models for studying the role of caveolae in integrin mechanosensing

To study the role of caveolae in integrin mechanosensing, we generated transgenic fibroblast lines by transducing Cav1KO MEFs with either Cav1- or PTRF-expressing retroviral vectors (Figure 1A-D).

**Figure 1.**
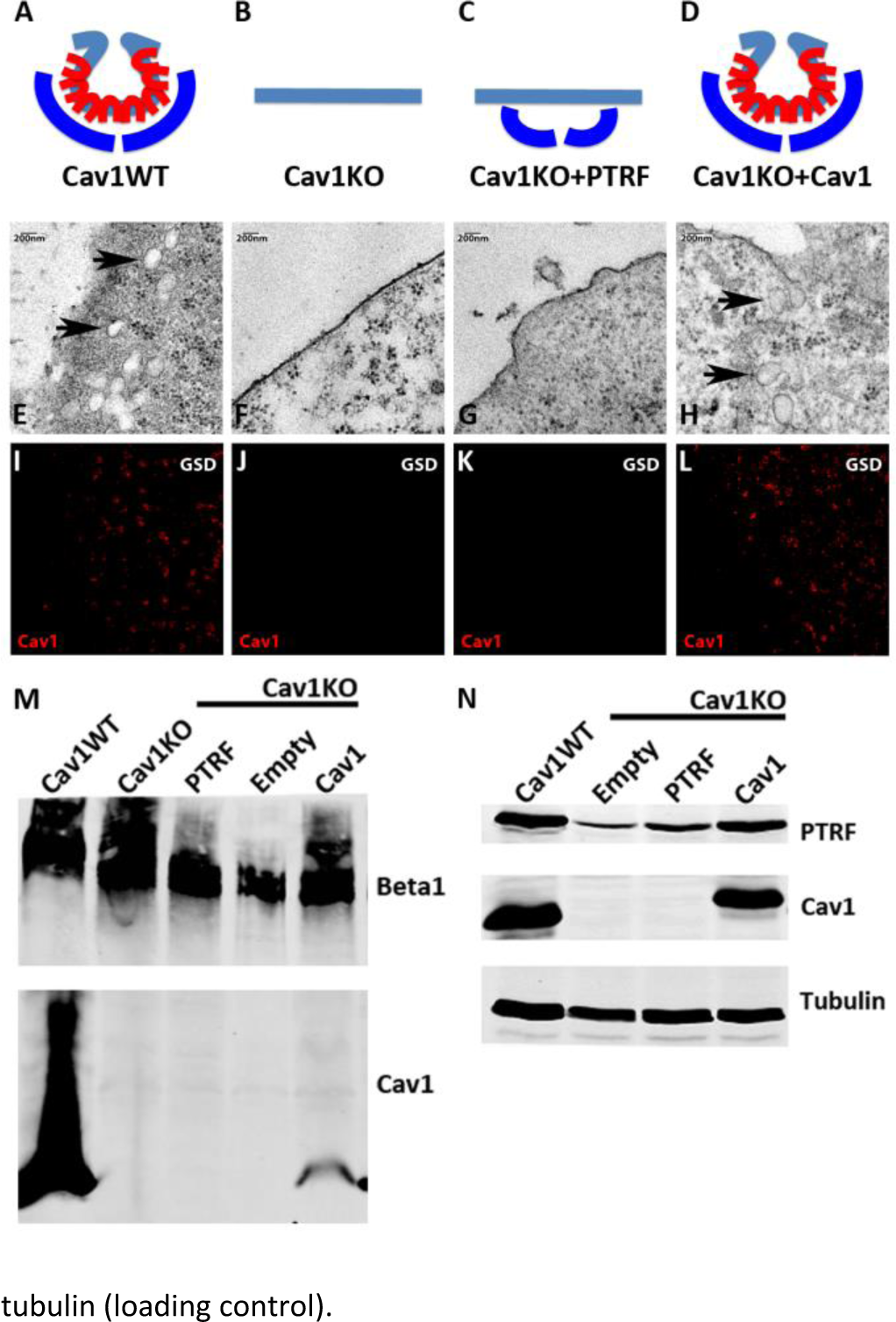
Caveolin-1-based genetic model characterization. (A-D) MEF caveolae-related phenotypes: PTRF is depicted in dark blue, Cav1 in red, and PM in light blue. (E-H) Electron microscopy images of MEF PM regions, showing the presence of caveolae only in (E) wild type MEFs and (H) Cav1-reconstituted Cav1KO MEFs (black arrows). (I-L) GSD-super-resolution MEF images. Cav1 is shown in red. (M) Biochemical PM fractionation of wild type and Cav1KO MEFs and Cav1KO MEFs reconstituted with PTRF, empty vector, and Cav1. Samples were immunoblotted for Cav1 and total β1-integrin 1 integrin (as both PM marker and loading control). (N) Western blot of total lysates from wild type, Cav1KO, and reconstituted Cav1 KO MEFs. Samples were immunoblotted for PTRF, Cav1, and tubulin (loading control). **Figure 1-source data 1-4** includes the full raw unedited blots (1 corresponding to Figure 1M, and 3 corresponding to Figure 1N) and the uncropped blots with the relevant bands labelled (2 corresponding to Figure 1M, and 4 corresponding to Figure 1N).

To validate the transgenic lines for the characterization of mechanosensing properties, we first analyzed the presence or absence of caveolae by electron microscopy (EM), detecting caveolae only in wild type and Cav1-reconstituted Cav1KO cells (Figure 1E-H, black arrows). Next, super-resolution microscopy analysis of Cav1 topographical distribution revealed that re-expressed Cav1 localizes to the PM and does not form aberrant aggregates (Figure 1I-L). Confirming the super-resolution imaging data, biochemical fractionation indicated that re-expressed Cav1 localizes to the PM (Figure 1M). Western blot analysis confirmed the expected Cav1 and PTRF expression in the different MEF lines and revealed near-endogenous expression levels in Cav1-reconstituted cells (Figure 1N, see also Figure 1-source data 1-4).

### Cav1 modulates integrin/cytoskeleton-dependent mechanosensing via Rho-independent mechanisms

To monitor the mechanosensing properties of the different transgenic lines, we used the MT technique. In this method, a pulsed magnetic force (1Hz, 1nN) is applied to pull magnetic beads attached to the cell surface (Figure 2A and 2B, and Suppl. Video 1). The magnetic beads oscillate in response to the pulsed force, and the local stiffness of the bead-cell adhesion can be measured as the ratio of the applied force to the bead movement. Cells able to detect the applied force respond through a phenomenon known as reinforcement, by which they progressively strengthen the cell-bead adhesion site, increasing its stiffness and thus reducing oscillation amplitude. Reinforcement can therefore be quantified as the change in adhesion stiffness over a specified time, providing a measure of cellular mechanosensing (Figure 2C and Suppl. Table 1; for more details on the technique, see^4^). We studied forces transmitted through integrins (adhesion strength) by coating beads with FN. As a control, we also studied forces transmitted non-specifically through the PM using beads coated with the sugar-binding lectin concanavalin A (ConA) (Figure 2D and Suppl. Video 2 and Video 3). We first assessed the tethering specificity of the two coatings by mixing FN-coated beads (prepared with non-labeled BSA) with ConA-coated beads labeled with Alexa 546-conjugated BSA (Figure 2E and 2F). Staining for phalloidin and 9EG7 antibody (which specifically recognizes β1-integrin in its active conformation^18^) revealed FN-coated beads surrounded by both signals, indicating engagement of both β1-integrin and the cell cytoskeleton; in contrast, ConA-coated beads were excluded from these stainings (Figure 2E and 2F) indicating they are only bound bulk PM, as previously reported ^19^.

**Figure 2.**
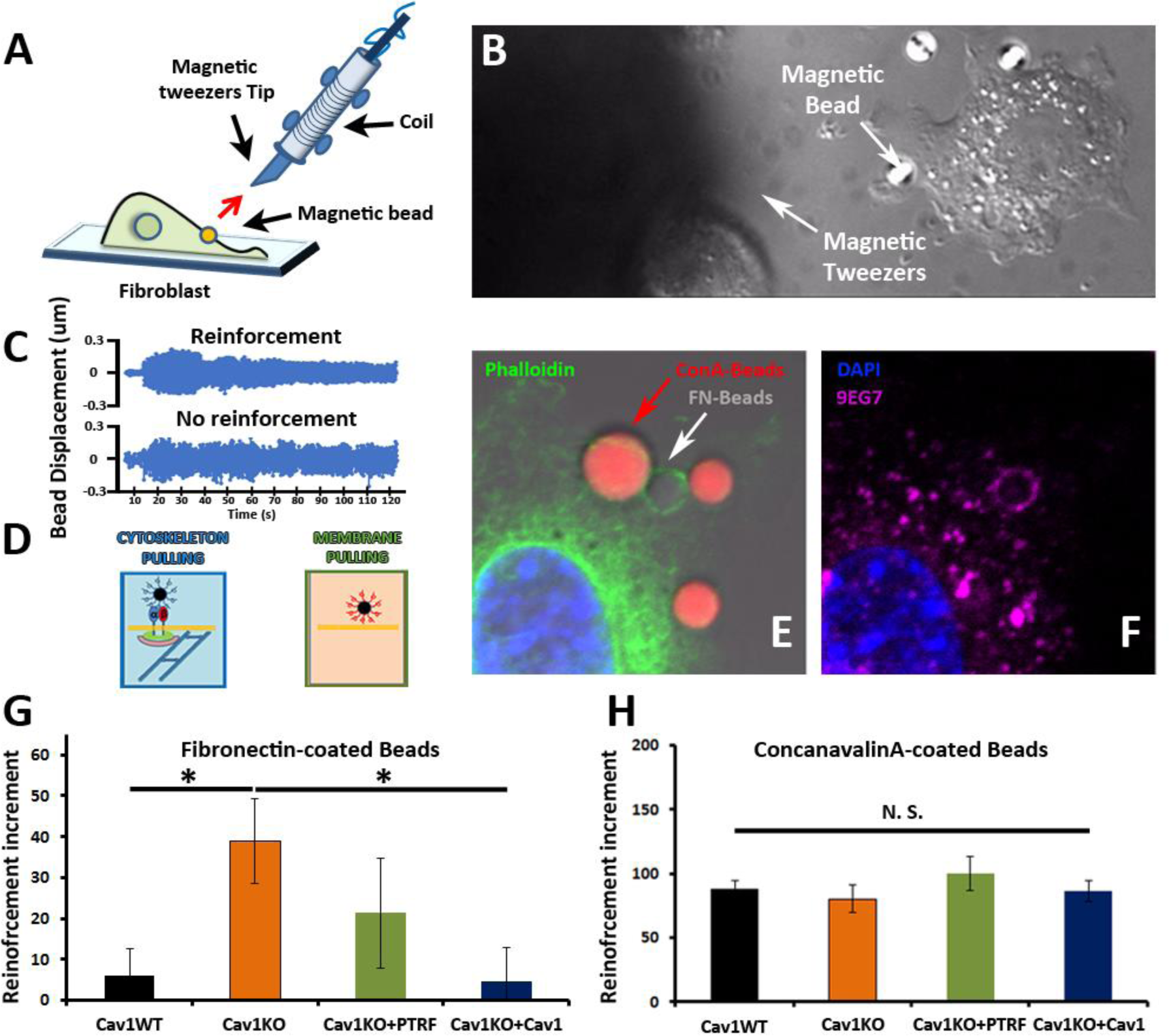
Cav1 KO MEFs show reinforced attachment to magnetic tweezers. (A) Reinforcement experiment scheme, indicating the fibroblast, the magnetic bead, and the magnetic tweezers apparatus. The red arrow represents the magnetic force exerted on the bead by the magnet. (B) Differential interference contrast (DIC) image showing a mouse embryonic fibroblast, the tip of the magnetic tweezers, and a magnetic bead (white arrows). (C) Examples of bead oscillation as a function of time in two conditions: with and without reinforcement. (D) Representation of the two magnetic beads coatings used: FN, which binds integrins, and ConA, which binds the bulk PM. (E and F) Confocal microscopy images showing a mouse embryonic fibroblast attached to concanavalin A-coated beads (red) and fibronectin-coated beads (gray). Actin staining is shown in green (phalloidin), active β1-integrin in magenta (9EG7 antibody), and DAPI in blue. Note how only fibronectin-coated beads present both phalloidin and 9EG7 staining. (G and H) Reinforcement increment (relative change in reinforcement over the entire experiment, calculated as the difference between the last and initial measurements) of different MEF genotypes for FN-coated beads (G) or ConA-coated beads (H); n ≥ 20 beads per genotype. Statistical comparisons were by *t*-test, with significance assigned at \**P*<0.05. N. S., non-significant. **See also Raw Data** Figures 2 and 3 which includes the raw data of experiments from Figure 2G and 2H.

We next analyzed the integrin/cytoskeleton-dependent mechanical response in different MEF lines. Cells were plated for 10 minutes on FN-coated coverslips. FN- or ConA-coated beads were then allowed to bind to the cell surface and subjected to magnetic pulses. Reinforcement was significantly higher in Cav1KO MEFs exposed to FN-coated beads than in wild type MEFs (Figure 2G). Reconstitution of Cav1KO MEFs with recombinant Cav1 rescued the WT phenotype, supporting that the Cav1KO phenotype is specific. Reconstitution of Cav1KO MEFs with PTRF induced a partial recovery that did not reach statistical significance. In contrast, MT pulling upon binding to ConA-coated beads yielded no significant differences across genotypes (Figure 2H). These results support an intrinsic role for Cav1/caveolae in determining PM mechanical properties, through integrin-dependent pathways.

Integrins link extracellular matrix (ECM) components, notably FN, to the cytoskeleton, transducing external forces into downstream effects and regulating responses such as cell contraction^3^. We previously showed that Cav1 regulates cell contraction by controlling Rho activity through the localization of p190RhoGAP^20^ within the PM^20, 21^. In the absence of Cav1, p190RhoGAP localization to liquid order domains increases, where it can bind and inhibit Rho activity^21^, thus attenuating cell contractility and associated ECM remodeling^20^.

This reduced cell contractility contrasts with the local stiffening response observed in our MT assays. Because the specific knock down of this p190RhoGAP isoform fully rescues other biomechanical phenotypes in Cav1KO MEFs^19, 20^, we assessed the impact of stably knocking down p190RhoGAP^21^ on the increased reinforcement observed in Cav1KO cells (Figure 3A-C and Suppl. Figure 1A, see also Figure 3-figure supplement 1-source data 1 and 2). While traction force microscopy confirmed higher overall contractility in p190RhoGAP-depleted Cav1KO MEFs as expected^21^ (Figure 3D), MT measurements revealed no decreased reinforcement upon p190RhoGAP knockdown (Figure 3C). These observations suggest that Cav1KO MEFs locally respond to FN-coated beads in a Rho-independent manner, as opposed to Rho-dependent whole cell contractility.

**Figure 3.**
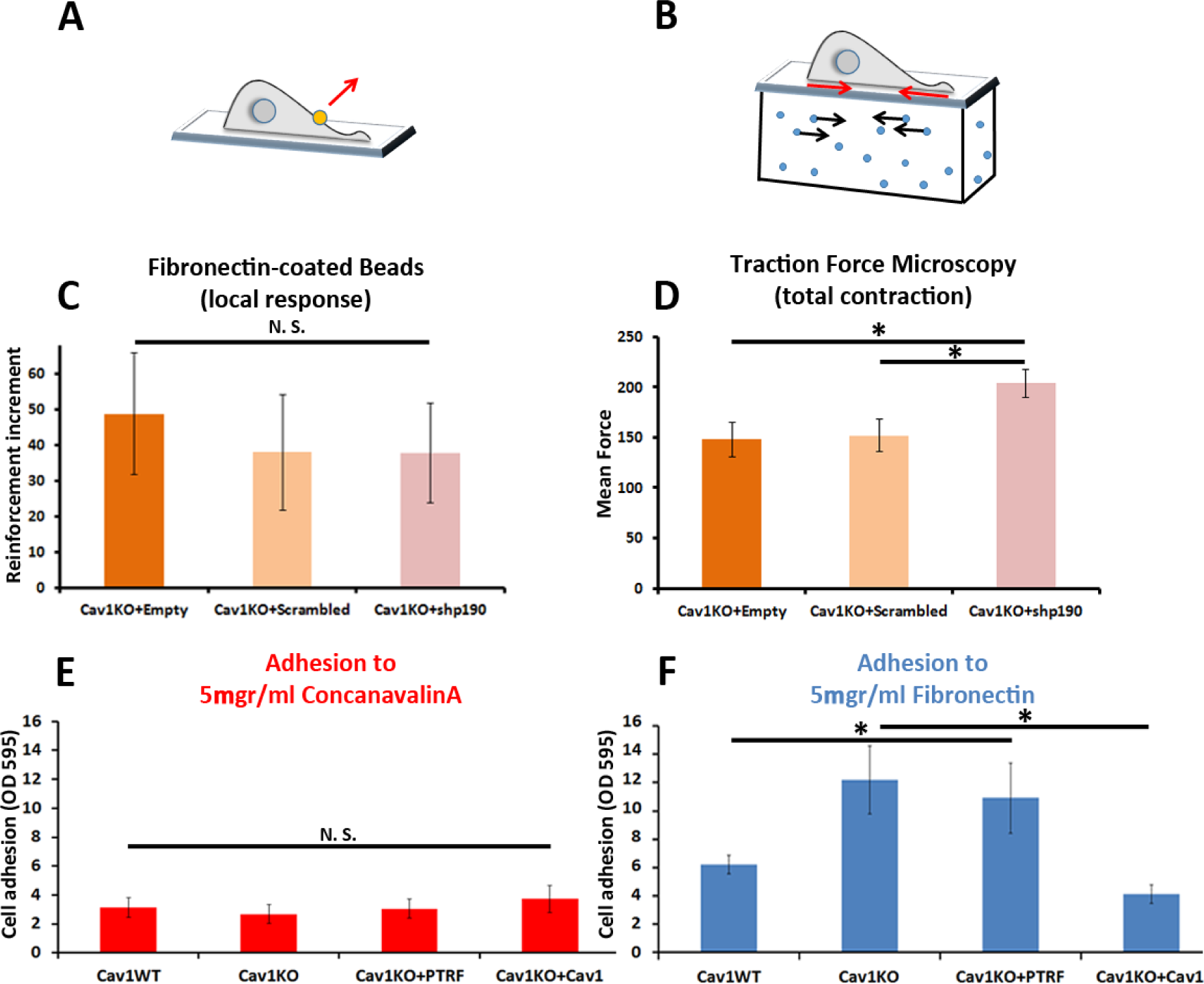
Cav1 KO MEFs show Rho-independent reinforcement and increased fibronectin adhesion. (A and B) Experimental schemes for (A) local force measurement (magnetic tweezers experiment, **reinforcement**) and (B) total force measurement (traction force microscopy, **total cell contraction**). (C) Effect of transfection with scrambled or shp190RhoGAP shRNA on the reinforcement increment in Cav1 KO MEFs (magnetic tweezers assay). Reinforcement increment refers to the relative change in reinforcement over the entire experiment, calculated as the difference between the last and initial measurements); n ≥ 20 beads per condition. (D) Effect of transfection with scrambled or shp190RhoGAP shRNA on mean total force contraction in Cav1 KO MEFs (traction force microscopy); n ≥ 12 cells per condition. (E and F) Relative adhesion of the indicated genotypes to plates coated with (E) 5ugr/ml ConA or (F) 5ugr/ml of FN. Measurements (absorbance, Optical density, OD, at 595nm from retained crystal violet dye, see material and methods) were normalized to values from adhesion to BSA-coated plates.; n ≥ 9 adhesion independent experiments. Statistical comparisons were by *t*-test, with significance assigned at \**P*<0.05. N. S., non-significant. **See also Raw Data** Figures 2 and 3 which includes the raw data of experiments from Figure 3C-3F.

### Loss of Cav1 increases fibronectin adhesion and active surface β1-integrin

To explore the role of cell adhesion in Cav1KO MEF reinforcement, we first analyzed the ability of the different MEF lines to adhere to FN- or ConA-coated plates. Whereas adhesion to ConA-coated plates did not differ between genotypes (Figure 3E), adhesion to FN was higher in Cav1KO MEFs and PTRF-reconstituted Cav1KO MEFs (Figure 3F). Cav1 re-expression rescued the wild type phenotype, supporting a specific role of Cav1 in cellular adhesion. However, PTRF reconstitution did not have a significant impact on Cav1KO MEFs adhesion, and was therefore not considered further in subsequent experiments. In additional experiments with different FN concentrations and other integrin-dependent coatings (collagen and vitronectin), Cav1KO MEFs showed always higher adhesion than wild type MEFs (Suppl. Figure 1B, 1C and 1D). The two major FN receptors are integrins α_5_β_1_ and α_v_β_3_; however, adhesion strength is mainly mediated by the clustering^4^ and activation^23, 24^ of integrin α_5_β_1_. To evaluate the role of β1-integrin in reinforcement, we imaged the active β1-integrin pool in Cav1KO and wild type MEFs by staining with 9EG7. The active β1-integrin signal was consistently stronger in Cav1KO MEFs than in wild type MEFs over a range of conditions: permeabilized and non-permeabilized cells, around FN-coated beads and at different spreading time-points (Suppl. Figure 1E-M). Cav1KO MEFs displayed stronger FN adhesion than wild type MEFs, a phenotype presumably derived from increased active β1-integrin at the cell surface.

### Cav1KO MEFs recycle β1-integrin faster than wild type MEFs

We first hypothesized that enhanced recruitment of active β1-integrin to FN beads in Cav1KO MEFs could be derived from increased lateral mobility. However, experiments measuring florescence recovery after photobleaching (FRAP) in Cav1KO and wild type MEFs expressing similar levels of GFP-fused β1-integrin (Figure 4A and 4B) revealed no significant differences in fluorescence recovery between the two cell populations, indicating that lateral mobility of β1-integrin was unaffected (Figure 4C).

**Figure 4.**
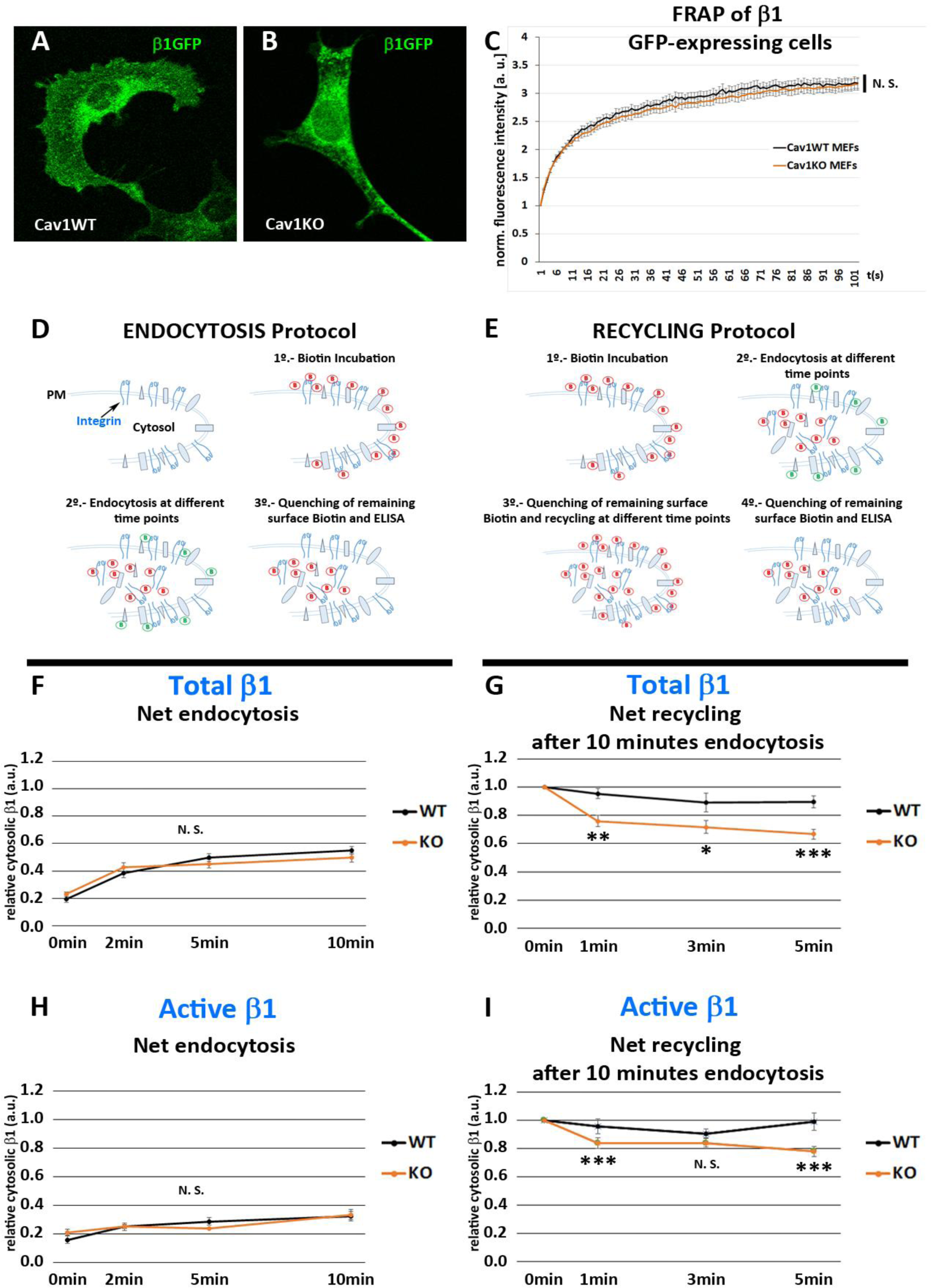
Cav1KO MEFs show faster β1-integrin recycling. (A and B) Confocal microscopy images of wild type and Cav1KO MEFs transfected with β1-integrin-GFP expression vector. (C) Normalized fluorescence intensity recovery after photo bleaching of wild type MEFs (black line) and Cav1KO MEFs (red line) at the indicated time points (the graph is representative of a minimum of 9 independent experiments in which between 8 and 10 cells were bleached per genotype). Statistical significance of between time-point differences was estimated by *t*-test; N. S., non-significant. (D and E) Experimental schemes for the analysis of endocytosis and recycling followed by ELISA (according to^25^). (F and H) Net endocytosis of (F) total β1 integrin and (H) active β1-integrin in wild type and Cav1KO MEFs at the time points indicated. Net endocytosis is expressed as internalized biotinylated β1 integrin (cytosolic) at each time point normalized to total biotinylated β1-integrin (internal and surface bound; see Materials and Methods); n ≥ 6 endocytosis assays per genotype. (G and I) Net recycling after 10 minutes endocytosis of (G) total and (I) active β1-integrin in wild type and Cav1KO MEFs at the time points indicated. Net recycling is expressed as internal biotinylated β1 integrin (cytosolic) at each time point normalized to time point 0 (which contains all the biotinylated β1 integrin internalized after 10 minutes of endocytosis, see Materials and Methods); then, decreasing values mean increased recycling; n = 10 recycling assays per genotype. Statistical comparisons were by *t*-test, with significance of between-group differences denoted \**P*<0.05, \*\**P*<0.01, or \*\*\**P*<0.001. N. S., non-significant. **See also Raw Data** Figures 4 and 5 which includes the raw data of experiments from Figure 4F-4I.

TIRF videos revealed that β1-integrins in wild type MEFs as stable, largely immobile structures; in contrast, β1-integrins in Cav1KO MEFs showed a dynamic behavior, rapidly appearing and disappearing from the PM plane (Suppl. Video 4 and Video 5 and Suppl. Figure 2A-B). Interestingly, integrins increased their dynamicity after treatment with high hypoosmotic pressure in wild type MEFs, mimicking Cav1KO phenotype (Suppl. Figure 2C-F and Suppl. Videos 6 and Video 7), suggesting that plasma membrane tension changes could affect integrin trafficking dynamics. In adherent cells, integrins undergo constant endocytic-exocytic shuttling to facilitate the dynamic regulation of cell adhesion^26^.

To study the effect of these dynamics on β1-integrin surface availability across our tested genotypes, we performed a series of endocytosis/recycling assays with an ELISA-based protocol^25^ (Figure 4D and 4E). We first analyzed the endocytic rates of total and active β1-integrin at early time-points (2, 5 and 10 minutes) during early spreading (2h after seeding), to recapitulate the conditions used in MT experiments (see Materials and Methods for details). Cav1KO and wild type MEFs showed no significant differences, indicating that β1 endocytosis is Cav1-independent at these early time points (Figure 4F and 4H). We next studied the recycling rates after allowing β1 endocytosis to proceed for 5 or 10 minutes; recycling was tested over two time-point sets: 2, 5, and 10 minutes and 1, 3, and 5 minutes. After 5 minutes of endocytosis, wild type and Cav1KO MEFs showed no major differences in total β1-integrin recycling rates at either time-point set (Suppl. Figure 2G and 2H). In contrast, after 10 minutes endocytosis, Cav1KO MEFs recycled total and active β1-integrin faster than wild type MEFs in the 1-3-5 minutes recycling set (Figure 4G and 4I). Importantly, reconstitution of Cav1KO MEFs with recombinant Cav1 rescued the WT phenotype, supporting that the Cav1KO phenotype is specific (Suppl. Figure 2I). These results suggest that β1-integrin is stabilized in the presence of Cav1 after 10 minutes endocytosis, whereas in Cav1KO MEFs its shuttling to PM is accelerated, increasing its surface availability and thus enhancing adhesion to FN-coated beads and reinforcement response. Different mechanisms could account for this stabilization, however, recent work in our lab showed increased exocytosis after loading wild type MEFs with cholesterol, phenocopying Cav1KO MEFs where cholesterol accrued in different endosomal compartments ^27^. To test if cholesterol could play a role in β1-integrin stabilization, we treated wild MEFs with either U18666A (which promotes cholesterol accumulation in endosomal compartments ^28^) or low-density lipoproteins, LDL (which also increases cholesterol content), and analyzed β1-integrin levels by ELISA. Interestingly, both treatments induced a significant increase in surface active β1-integrin (Suppl. Figure 2J), that was also accompanied by an increased colocalization with EEA-1 positive endosomes in LDL-treated cells (Suppl Figure 2K-M). These results might be indicative of a Cav1-dependent cholesterol threshold above which β1-integrin trafficking is altered, phenocopying Cav1KO MEFs where both surface and intracellular active β1-integrin levels are increased.

### CLIC/GEEC-dependent uptake contributes to β1-integrin endocytosis in Cav1KO MEFs

β1-integrin endocytosis has been suggested to be Cav1-dependent in different systems^29, 30^; however, we observed no differences between Cav1KO and wild type MEFs in net β1-integrin endocytosis. This discrepancy might reflect assay timings, as we studied early time points (2, 5, and 10 minutes), whereas previous studies focused on endocytosis over longer time periods. To investigate how Cav1KO MEFs achieve similar levels of β1-integrin endocytosis as wild type MEFs, we decided to analyze endocytosis via Clathrin independent carriers (CLIC), especially because this endocytic modality is negatively regulated by caveolar proteins, including Cav1 and PTRF^31, 32^. Furthermore, this endocytic route is highly relevant for fibroblast trafficking dynamics—accounting for ∼3-fold internalized volume as compared to clathrin-mediated endocytosis—and β1-integrin is a specific cargo^33^. Interestingly, cells lacking Cav1 increase CLIC endocytosis through the small GTPase Cdc42 activation^32^, which is known to regulate this endocytic pathway^34^. We therefore asked ourselves whether Cav1KO MEFs might preferentially endocytose β1-integrin via this mechanism. We treated wild type and Cav1KO MEFs with the Cdc42 inhibitor ML141^35^, which inhibits the CLIC-GEEC (<u>cl</u>athrin independent <u>c</u>arrier/<u>G</u>PI <u>e</u>nriched <u>e</u>ndocytic <u>c</u>ompartment) pathway of endocytosis, involved in fluid-phase uptake and the entry of many specific cargoes, including integrins^33, 36^. ML141 significantly reduced β1-integrin endocytosis in Cav1KO MEFs, whereas wild type MEFs were unaffected (Figure 5A and 5B), indicating that β1-integrin is partially endocytosed through the CLIC/GEEC pathway in Cav1KO MEFs.

**Figure 5.**
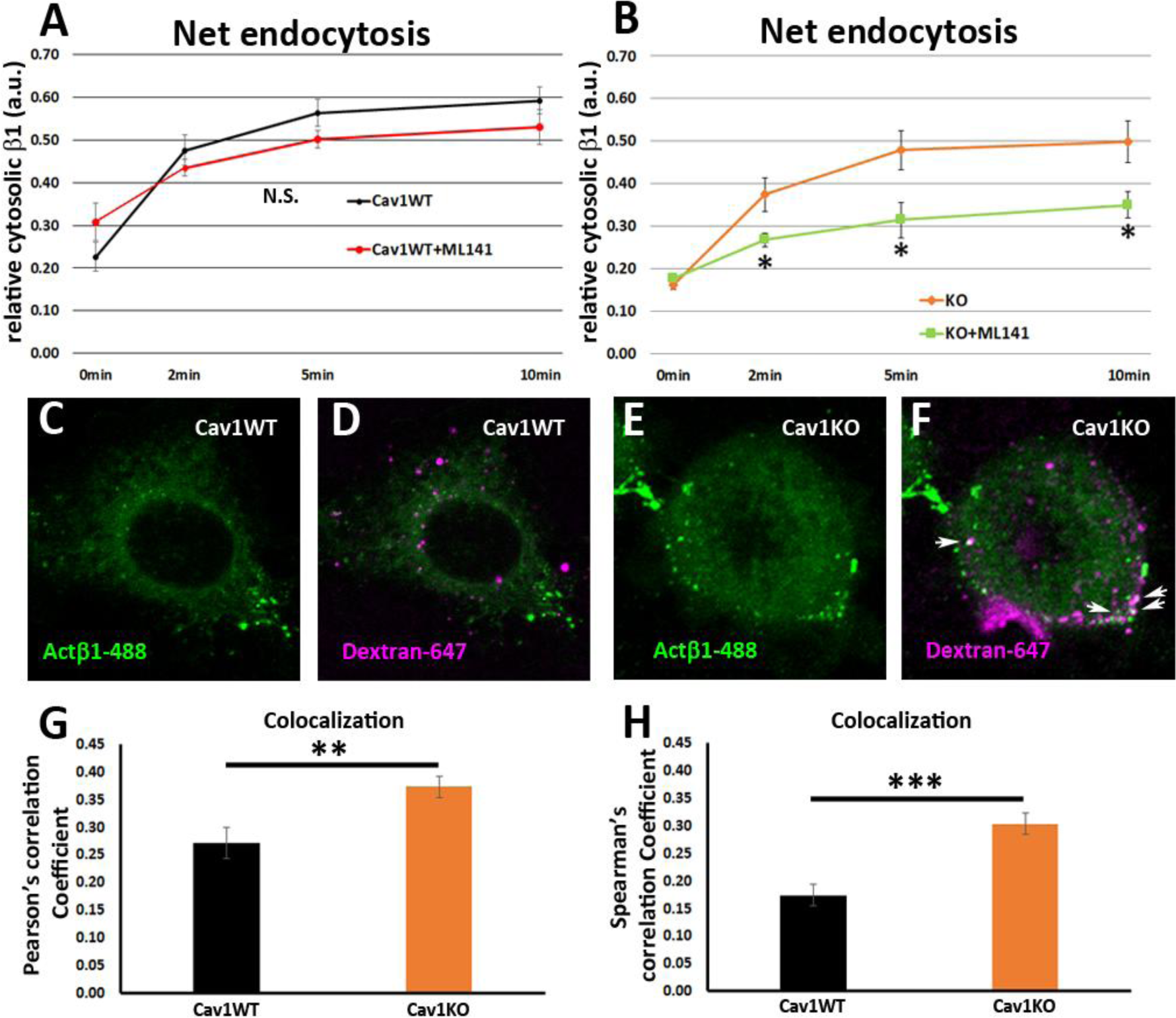
Cav1KO MEFs take up β1-integrin by CLIC endocytosis. (A and B) Net endocytosis (normalized to Total biotinylated β1-integrin) at the time points indicated. (A) wild type MEFs treated with ML141 (red line) and untreated controls (black line). (B) Cav1KO MEFs treated with ML141 (green line) and untreated controls (orange line); n ≥ 5 endocytosis assays per genotype. (C-F) Confocal microscopy images of wild type MEFs (C and D) and Cav1KO MEFs (E and F) incubated with anti-active β1-Alexa 488 antibody (green) for 1 hour at 4°C followed by incubation with dextran-Alexa 647 (magenta) for 3 minutes at 37°C. White arrows in F mark colocalization between β1-488– positive particles and dextran-647–positive particles in Cav1KO MEFs. (G and H) Quantification of colocalization between active β1-Alexa 488 and dextran-647, expressed as Pearson’s correlation coefficient (G) or Spearman’s correlation coefficient (H); n ≥ 18 cells per genotype. Statistical significance of differences across indicated conditions was assessed by *t*-test: * *p*<0.05 P\*\**p*<0.01; \*\*\**p*<0.001 N. S., non-significant. **See also Raw Data** Figures 4 and 5 which includes the raw data of experiments from Figure 5A, 5B, 5G and 5H.

In accordance with this finding, Cav1KO MEFs displayed significantly higher uptake of the fluid-phase endocytosis marker dextran^37^, as well as elevated colocalization of dextran and Alexa-488-labeled β1-integrin (Figure 5C-5H and Suppl. Figure 3A-C), as compared to wild type cells. Thus, in the absence of Cav1, early endocytosis of β1-integrin occurs at least in part through CLIC uptake, which provides an alternative entry route that would compensate for lack of Cav1-dependent internalization. To further delineate the relative contribution of the different endocytic routes to beta1 integrin endocytosis, we performed a series of co-localization studies of active β1-integrin and previously characterized markers for caveolar-dependent (BODIPY-LacCer), CLIC-dependent (CD44) and clathrin-dependent (transferrin, Tnf-568) endocytosis ^38–42^. Consistent with previous results, β1-integrin endocytosis was mainly Cav1-dependent in wild type MEFs as it: i) co-localized with Cav1 and LacCer, ii) was significantly reduced upon genistein treatment (a caveolar endocytosis inhibitor^43^) and iii) was unaffected by ML141 treatment (the CLIC inhibitor) (Suppl. Figure 3D-K). On the other hand, β1-integrin endocytosis was mainly CLIC-dependent in Cav1KO MEFs as it: i) co-localized with CD44 and ii) was significantly reduced upon ML141 treatment as compared to wild type MEFs (Suppl. Figure 3L-P). Finally, no significant differences were found in clathrin-dependent beta 1 integrin endocytosis between wild type and Cav1KO MEFs (Suppl. Figure 3Q-S). Altogether, these results further prove that β1-integrin endocytosis is mainly endocytosed by CLIC-dependent mechanisms in Cav1KO MEFs.

### Cav1 is required for Rab11-dependent recycling of β1-integrin

β1-integrin follows the canonical Rab21-Rab11-dependent endosomal trafficking route— which takes longer times to recycle back to the PM—in wild type MEFs; while other paralogs, such as, for instance, β3-integrin follow a Rab4-dependent “short” loop^44–46^. As stated above, this scenario is in accordance with integrin β1 being localized to Rab11-positive endosomal comparments after 10 minutes endocytosis in wild type MEFs. In contrast, in the absence of Cav1, β1-integrin is partially sorted to a CLIC-dependent endosomal compartment in Cav1KO MEFs, from which it might be recycled to the PM following different dynamics. We have previously reported that integrins are rapidly delivered to nascent focal contacts in the absence of Cav1^20^. The recycling of CLIC cargo proteins is controlled by a number of factors including several Rabs such as Rab22a^47^, which has been shown to collaborate with the microtubule and tethering protein HOOK1 during this process^48^. Interestingly, knocking down HOOK1 shifts the trafficking of CLIC cargo proteins from recycling to endosomal targeting, such as the surface glicoprotein CD147, which accumulates in the early endosomal antigen-1 positive compartment (EEA-1)^48^. To analyze whether HOOK1 is required for active β1−integrin recycling, we transfected Cav1KO MEFs with siRNA against HOOK1 and studied the colocalization of EEA1 and 9EG7 (i.e. active β1−integrin) immunolabelling. Whereas HOOK1-deficient Cav1KO MEFs showed increased CD147 and EEA1 colocalization as expected^48^, no significant differences were observed for β1 integrin (Figure 6A-6E). Accordingly, surface active β1−integrin showed similar levels in both HOOK1-deficient and control Cav1KO MEFs, even 72h after siRNA treatment (Suppl. Figure 4A). These results indicate that HOOK1 is not required for β1−integrin recycling in Cav1KO MEFs. We then decided to analyze the contribution of canonical Rab11 and Rab4-dependent recycling routes, using dominant-negative (DN) mutants described previously: Rab11 N124I, and Rab4 S22N, respectively^45, 49^. We first assessed the impact of disrupting ‘long loop’-dependent recycling upon expression of the Rab11 DN mutant^49^, as assessed by the degree of colocalization between the endosomal compartment (LBPA, lysobisphosphatidic acid, a late endosomal marker) and active β1−integrin (9EG7 label) in either wild type or Cav1KO cells. While we observed significant differences in the colocalization of 9EG7 and LBPA labels when comparing Cav1KO cells transfected with the Rab11 DN mutant, to non-transfected Cav1KO cells, no significant differences were observed on the surface exposure of active β1−integrin in the same cells (Figure 6G, 6I, 6J and Suppl. Figure 4B). In contrast, wild type MEFs showed increased colocalization between 9EG7 and LBPA-positive vesicles (Figure 6F, 6H and 6J) and reduced surface active β1−integrin levels upon Rab11 DN transfection (Suppl. Figure 4B), suggesting that this is the main recycling pathway in wild type cells. We then assessed the impact of expressing a Rab4 S22N DN mutant (which blocks a ‘short loop’-dependent recycling^45^) on the trafficking of active β1−integrin in either wild type or Cav1KO cells. Whereas no significant differences in 9EG7-EEA1 colocalization was observed upon disrupting Rab4-dependent trafficking in wild type MEFs, expression of the Rab4 DN mutant increased the colocalization between both labels in Cav1KO cells (Figure 6K-6O). This was consistent with a significant reduction in surface active β1−integrin levels in Cav1KO cells expressing the Rab4 DN mutant, as compared to non-transfected Cav1KO cells, while no difference was observed for wild type cells (Suppl. Figure 4C). Taken together, these results suggest that in the absence of Cav1, β1−integrin recycling is partially switched from “slow” Rab11-dependent, to “fast” Rab4-dependent, recycling.

**Figure 6.**
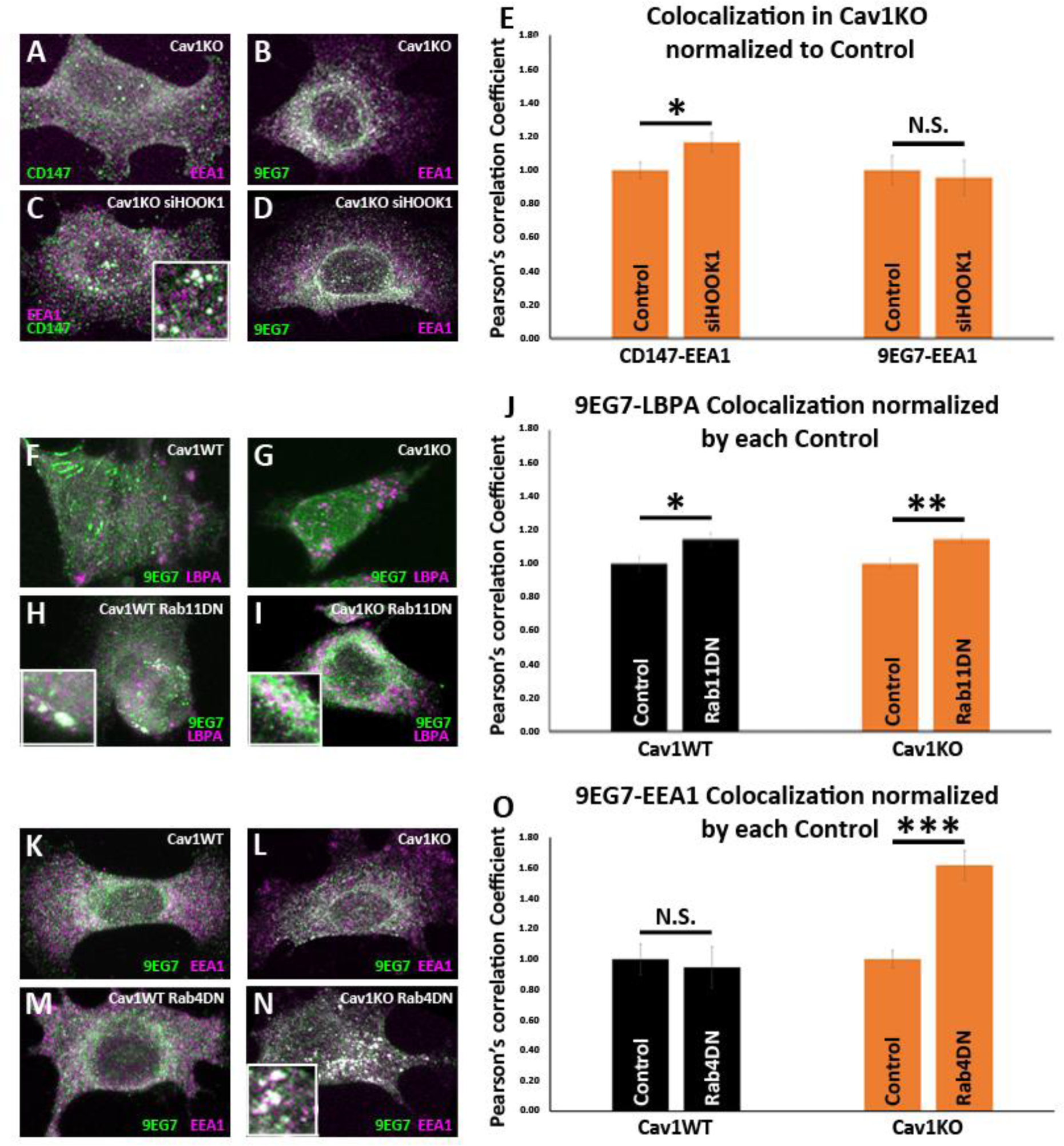
Cav1 is required for β1-integrin Rab11-dependent recycling. (A-D) Confocal microscopy images of Cav1KO MEFs stained for CD147 (green in A and C), active β1-integrin (9EG7 antibody, green in B and D) and EEA1 (magenta), either non-treated (A and B) or treated with siRNA against HOOK1 for 48h (C and D). (E) Quantification of colocalization between EEA1 and CD147 or 9EG7 normalized to control, expressed as Pearson’s correlation coefficient; n ≥ 20 cells per condition. (F-I) Confocal microscopy images of wild type (F and H) or Cav1KO MEFs (G and I), stained for active β1-integrin (9EG7 antibody, green) and LBPA (magenta), either non-transfected (F and G) or transfected with Rab11 N124I dominant negative mutant for 48h (H and I). (J) Quantification of colocalization between LBPA and 9EG7 normalized by each control, expressed as Pearson’s correlation coefficient; n ≥ 20 cells per condition. (K-N) Confocal microscopy images of wild type (K and M) or Cav1KO MEFs (L and N), stained for active β1-integrin (9EG7 antibody, green) and EEA-1 (magenta), either non-transfected (K and L) or transfected with Rab4 S22N dominant negative mutant for 48h (M and N). (O) Quantification of colocalization between EEA-1 and 9EG7 normalized to each control, expressed as Pearson’s correlation coefficient; n ≥ 30 cells per condition. Colocalization was analyzed using the plugin Intensity Correlation Analysis (Fiji^50^). Statistical significance of differences across indicated conditions was assessed by *t*-test: * *p*<0.05; P\*\**p*<0.01; \*\*\**p*<0.001 N. S., non-significant. **See also Raw Data** Figures 6 and 7 which includes the raw data of experiments from Figure 6E, 6J and 6O.

### Hypoosmotic shock increases β1-integrin activation in Cav1KO MEFs

Many studies have shown that membrane trafficking and membrane tension are tightly coupled^51, 52^. For example, increases in membrane tension increase exocytosis from the endocytic recycling compartment^53^. Cav1KO MEFs lacking caveolae cannot buffer membrane tension properly^9^. This can lead to increased exocytosis^54^, as we observed in the specific case of β1-integrin recycling, which in turn increases adhesion to FN-coated beads. However, PM tension can also affect adhesion more directly through increasing integrin activation independently from the cytoskeleton^55^ and in a ligand-independent mechanism^7^. To study the role of caveolae in membrane-tension–induced β1-integrin activation, we exposed cells to hypoosmotic conditions. Cav1KO and wild type MEFs were incubated for 10 minutes in DMEM diluted 1:10 in distilled water, fixed, and stained for 9EG7. Hypoosmotic shock sharply increased β1-integrin activation in Cav1KO MEFs, whereas no significant change was observed in wild type MEFs (Figure 7A-7F; similar results were obtained with hypoosmotic shock exposure for 30 seconds and 1 minute; data not shown). We also studied the amount of active β1-integrin around FN-coated beads before and after magnetic twisting (Suppl. Video 8), observing a significant tension-induced increase in Cav1KO MEFs (Suppl. Figure 5A-5J). We confirmed this phenotype is caveolae-dependent, Cav1-independent, as PTRFKO MEFs (that lack caveolae but still express Cav1^13^) also show increased β1-integrin activation upon hypoosmotic shock (Suppl. Figure 5K-5M). Strikingly, β1-integrin activation was also observed in Cav1WT MEFs after both longer hypoosmotic treatments and with higher hypoosmotic pressures (Suppl. Figure 5N-5W). This might indicate that caveolae restrict integrin activation upon changes in plasma membrane tension until their buffering capacity is exhausted, and therefore Cav1KO MEFs lack the ability to adapt integrin activation to mechanical stress.

**Figure 7.**
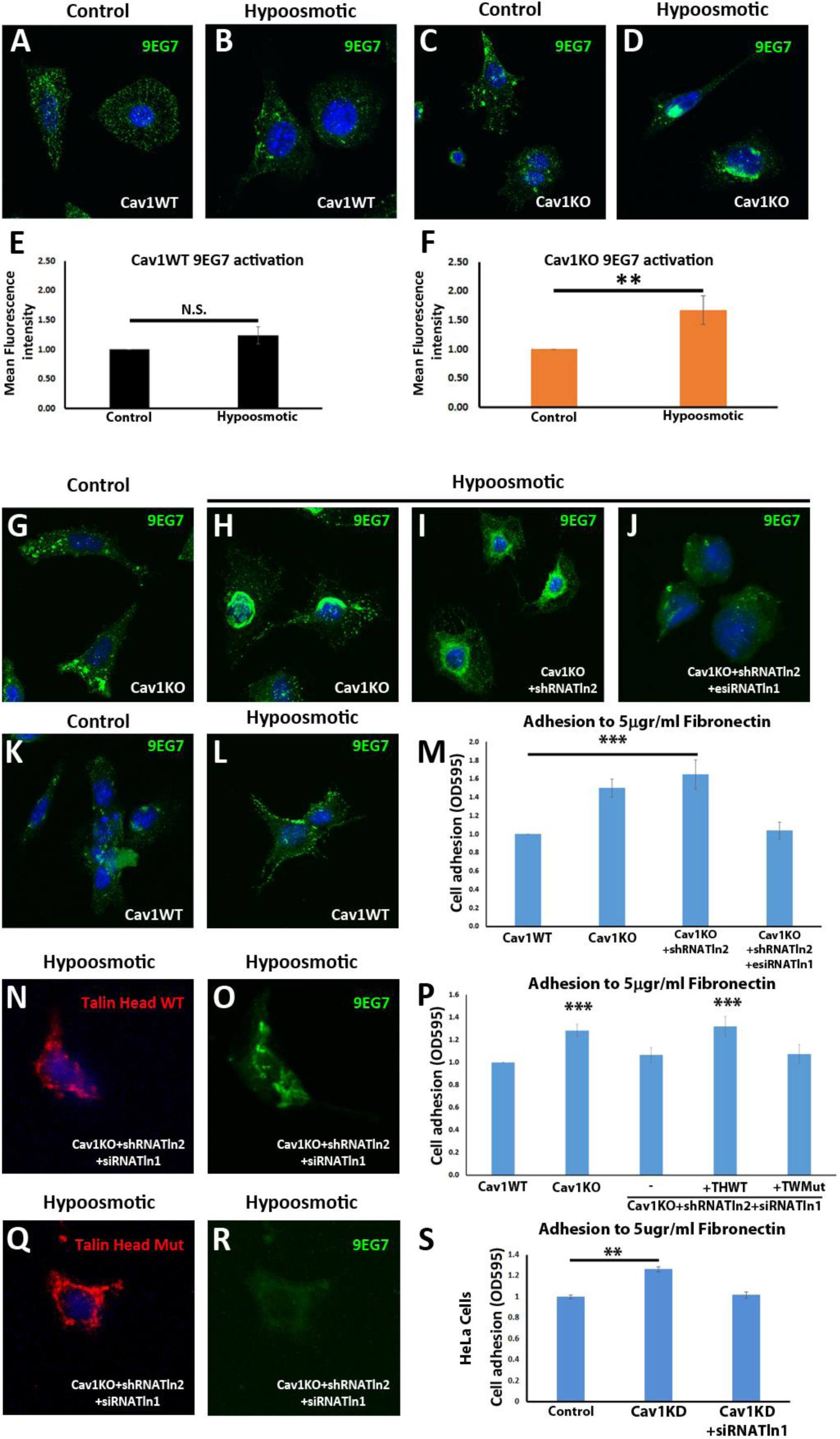
Talin is required for the enhanced adhesion and β1-integrin activation phenotype of Cav1KO MEFs. (A-D) Confocal microscopy images of wild type MEFs (A and B) and Cav1KO MEFs (C and D) stained for active β1-integrin (9EG7 antibody, green) after culture in standard medium (control in A and C) or hypoosmotic medium (diluted 1:10; B and D) for 10 minutes at 37°C. DAPI is shown in blue. (E and F) 9EG7 mean fluorescence intensity in control and hypoosmotic shock-exposed wild type (E) or Cav1KO MEFs (F); n ≥ 35 cells per genotype. (G-J) Confocal microscopy images of active β1-integrin staining (9EG7 antibody, green) in Cav1KO MEFs subject to indicated RNAi treatments and cultured for 10 minutes at 37°C in standard medium (control) or hypoosmotic medium (diluted 1:10). (K and L) Active β1-integrin immunostaining in wild type MEFs cultured in standard (control) or hypoosmotic medium. DAPI counterstain is shown in blue. (M, P and S) Relative adhesion of MEFs of the indicated genotypes (M and P) or HeLa Cells (S, wild type – control -; knocking down Cav1-Cav1KD-; and knocking down both Cav1 and Talin1 -Cav1KD+siTnl1) to plates coated with 5 µg/ml FN; n ≥ 18 cells in 3 independent adhesion experiments. THWT: Talin Head wild type; THMut: Tlain Head Mutant. (N, O, Q and R) Confocal microscopy images of talin head domain (N, Q) and active β1-integrin (9EG7 antibody; O, R) in Cav1KO MEFs transfected with Tln2 shRNA and Tln1 siRNA plus either WT Talin Head (N and O) or Mutant Talin Head (Q and R). Before immunostaining, cells were cultured for 10 minutes at 37°C in hypoosmotic medium (diluted 1:10). DAPI is shown in blue. All immunostainings in this figure were performed following the extracellular staining (see material and methods for more details). Statistical significance of differences across indicated conditions was assessed by *t*-test: * *p*<0.05 P\*\**p*<0.01; \*\*\**p*<0.001 N. S., non-significant. **See also Raw Data** Figures 6 and 7 which includes the raw data of experiments from Figure 7M, 7P and 7S, and **Raw Data Suppl. Figures 5 and 6** which includes the raw data of experiments from Figure 7E and 7F pooled together in Suppl. Figure 5V and 5W.

### Talin supports increased β1-integrin activation and recycling in Cav1KO MEFs

Integrin activation is controlled by a number of intracellular proteins^6^. One of them, talin, regulates integrin adhesion strength by interacting through its amino-terminal FERM domain (the talin head domain)^56^. This interaction is necessary and sufficient to induce inside-out integrin activation^57, 58^. To study the possible role of talin in β1-integrin activation in Cav1KO MEFs, we first confirmed β1-integrin binding competence after hypoosmotic treatment. Co-localization between β1-integrin and both soluble fibronectin and talin proved its active conformation (Suppl. Figure 6B-6E). We then analyzed the effect on adhesion and activation after silencing Tln1 and Tln2. Interestingly, Tln2-silencing did not significantly affect either hypoosmotic-induced β1-integrin activation or adhesion in Cav1KO MEFs (Figure 7G-7I, 7K-7M and Suppl. Figure 6A). In contrast, simultaneous silencing of Tln1 and Tln2 in Cav1KO MEFs reduced hypoosmotic-induced β1 activation and reduced adhesion, rescuing the wild type phenotype (Figure 7J, 7M and Suppl. Figure 6A, see also Figure 7-figure supplement 6-source data 1 and 2). The same result was observed in Cav1-silenced HeLa cells (Figure 7S). Surprisingly, Tln1/2 knockdown also affected integrin trafficking, as surface active β1-integrin was significantly reduced and consistently increased intracellularly in EEA-1 positive endosomes (Suppl. Figure 6F-6M). Given that the talin head domain binds and activates integrins^40, 58^, we studied the ability of the wild type talin head to rescue adhesion in Tln1- and Tln2-silenced Cav1KO MEFs. As a control, we transfected cells with a dominant-negative talin-head mutant (L325R) that does not activate integrins nor link them to the cytoskeleton^59^. Expression of the WT talin head, but not the L325R mutant, rescued both the activation and the adhesion ability of Cav1KO MEFs (Figure 7N-7P, 7Q and 7R). The talin head thus regulates adhesion and β1-integrin activation in Cav1KO MEFs.

## Discussion

Cellular mechanosensing is dependent on both integrins and caveolae, but how these two are coupled in this cellular response is poorly understood, especially during early steps of mechanoadaptation. Our MT-pulling experiment results show that Cav1KO MEFs adhere more strongly than wild type MEFs to FN-coated beads. This is consistent with the higher adhesion behavior of Cav1KO MEFs as compared to wild type MEFs in substrate adhesion assays. Fibronectin binds to integrins, and β1 is its main receptor. The analysis of β1 distribution revealed an increased surface pool of active β1-integrin in Cav1KO MEFs. TIRF imaging and FRAP measurements indicated that β1-integrin has a more dynamic behavior in Cav1KO MEFs, appearing at and disappearing from the membrane plane faster than in wild type MEFs, through mechanisms distinct from diffusion. This prompted us to study endocytosis/recycling rates. ELISA-based assays revealed no differences between Cav1KO and wild type MEFs in endocytosis rate at the time points analyzed, suggesting that Cav1 does not play a specific role. This finding contrasts with other reports showing that Cav1 is required for proper β1-integrin endocytosis^26, 30^; however, these observations were not made in a Cav1KO background and were performed at later time points than examined in our experiments (as we wanted to analyze specifically early integrin mechanosensing events, studied in the MT experiments). This time difference suggests that net integrin endocytosis at early time points, could be compensated by other mechanisms in Cav1KO MEFs. Furthermore, and in accordance with previous reports^60^, Cav1 could be restricting the endocytosis of a pool of β1-integrin in wild type MEFs (where it is mainly endocytosed through caveolae-dependent mechanisms), and becomes freed in Cav1KO MEFs to follow a different entry route. Indeed, our results indicate that β1-integrin is partially taken up in Cav1KO MEFs through CLIC/GEEC endocytosis, which provides a fast entry route, as reported for other cargoes^31, 32, 61^. Interestingly, net endocytosis of active β1-integrins is higher than that of inactive pools^62, 63^. This is consistent with our data showing that Cav1KO MEFs display higher 9EG7 signal inside the cell, and harmonizes with our findings that Cav1 genetic deficiency enhances CLIC endocytosis.

Differences between Cav1KO and wild type MEFs were apparent when we analyzed β1-integrin recycling rates at early time points after 10 minutes of endocytosis, being faster in Cav1KO MEFs. Interestingly, no differences in recycling were observed after 5 minutes of endocytosis, suggesting that in the presence of Cav1, β1-integrin becomes stabilized over time. This could potentially depend on cholesterol levels as loading wild type MEFs with cholesterol increases both surface and endosomal active β1-integrin availability, phenocopying Cav1KO MEFs phenotype. Raising endosomal cholesterol levels leads to increased exocytic activity in wild type MEFs, mirroring Cav1KO MEFs cholesterol accumulation, as work in our lab has previously demonstrated^27^. This suggests that there might be a cholesterol threshold above which β1-integrin trafficking is dysfunctional, as it happens in the absence of Cav1. Furthermore, our results suggest that Cav1 is required for Rab11-dependent recycling of β1-integrin, which takes longer to reach the PM. Interestingly, in the absence of Cav1, β1-integrin accumulates preferentially at EEA-1-positive vesicles, following the Rab4-dependent “short” recycling loop. This is consistent with our previous observations of Cav1 playing a role in determining the migration mode of fibroblasts^20^, alternating between persistent or random migration. Faster recycling in Cav1KO MEFs can account for the elevated β1-integrin surface availability and therefore explain the resulting reinforcement. This interpretation is further supported by our previous findings that Cav1KO MEFs have a higher number of small focal adhesions that are rapidly turned over^20^, since short but frequent integrin-ECM contacts would strengthen adhesion over time (ECM-coated beads in the MT experimental design). In the same report, we showed that Cav1KO MEFs have low Rho activity resulting in impaired actomyosin contraction^20^, a condition that alters the overall cell mechanical response, as revealed in our traction force experiments. However, in the present study, Rho appears to be dispensable for observed differences in reinforcement. Consistently, previous reports have shown initial integrin adhesion in the absence of cytoskeleton connection, suggesting that early mechanosensing could be locally triggered^5, 64, 65^. This actomyosin-independent adhesion can derive from increased PM tension^55^, which has also been shown to induce ligand-independent integrin activation^7^. Cav1KO MEFs lack caveolae and are unable to buffer membrane tension upon mechanical stress^9^. In this condition, integrin activation at a lower force threshold might be facilitated by an easier switch of β1-integrin to its active conformation, as we observed upon hypoosmotic treatment. This increased sensitivity to membrane tension contributes, together with increased recycling, to the higher surface availability of active β1-integrin we observed in Cav1KO MEFs. Interestingly, we have also observed integrin activation in Cav1WT MEFs but after longer treatment times and with increased hypoosmotic pressure. These results suggest that caveolae membrane buffering is limiting integrin activation before a plasma membrane tension threshold is reached. Previous studies have shown that talin controls inside-out integrin activation^66, 67^ through its head domain^57, 58^. Interestingly, our results show that talin expression is required both for adhesion and for integrin activation in Cav1KO MEFs. It also seems to play a role in integrin recycling, as Tln1/2 knockdown altered integrin trafficking. Importantly, depletion of both talin paralogs does not affect initial cell spreading^58^. This is consistent with our observation that talin depletion does not affect initial attachment of Cav1KO MEFs to the substrate (Figure 7M and 7P, see material and methods for details), as cells were allowed in our adhesion assays to spread for 30min prior to measurements. In contrast, cells were allowed to spread and form attachments for at least 48 hours after reverse siRNA transfection in experiments were hypoosmotic shock was induced, and—while still attached—Cav1KO/KD cells depleted for Talins were rounder and less spread (compare Figure 7H with 7J) than control cells. Rescue experiments indicated that this situation is mainly supported by the talin head domain. This suggests the intriguing possibility that membrane tension could favor local integrin–talin-head interaction without force transmission in Cav1KO MEFs. These observations suggest that the cellular cytoskeleton might play only a minor role in the early cellular response to mechanical stress. A similar conclusion was recently prompted by the detection of mechanical-stress–induced integrin recruitment in the absence of significant cytoskeletal changes^5^.

Our results indicate that caveolae impact both integrin surface availability, through adjusting recycling, and integrin activation, through membrane tension regulation and talin activity. Cav1KO MEFs, lacking this control mechanism, show both increased β1-integrin recycling and surface activation. This results in dysregulated early mechanosensing (Figure 8) and subsequent inability to properly sense environmental stiffness, a situation known to impact tumorigenesis^68^ and stem cell differentiation^25^. Mechanobiology is an emerging field, essential to understand how cells and tissues adapt to their environment in health and disease^69^. Our study provides novel insight on the role of caveolae in early integrin mechanosensing, revealing a new layer of complexity at the interface of physics and biology.

**Figure 8.**
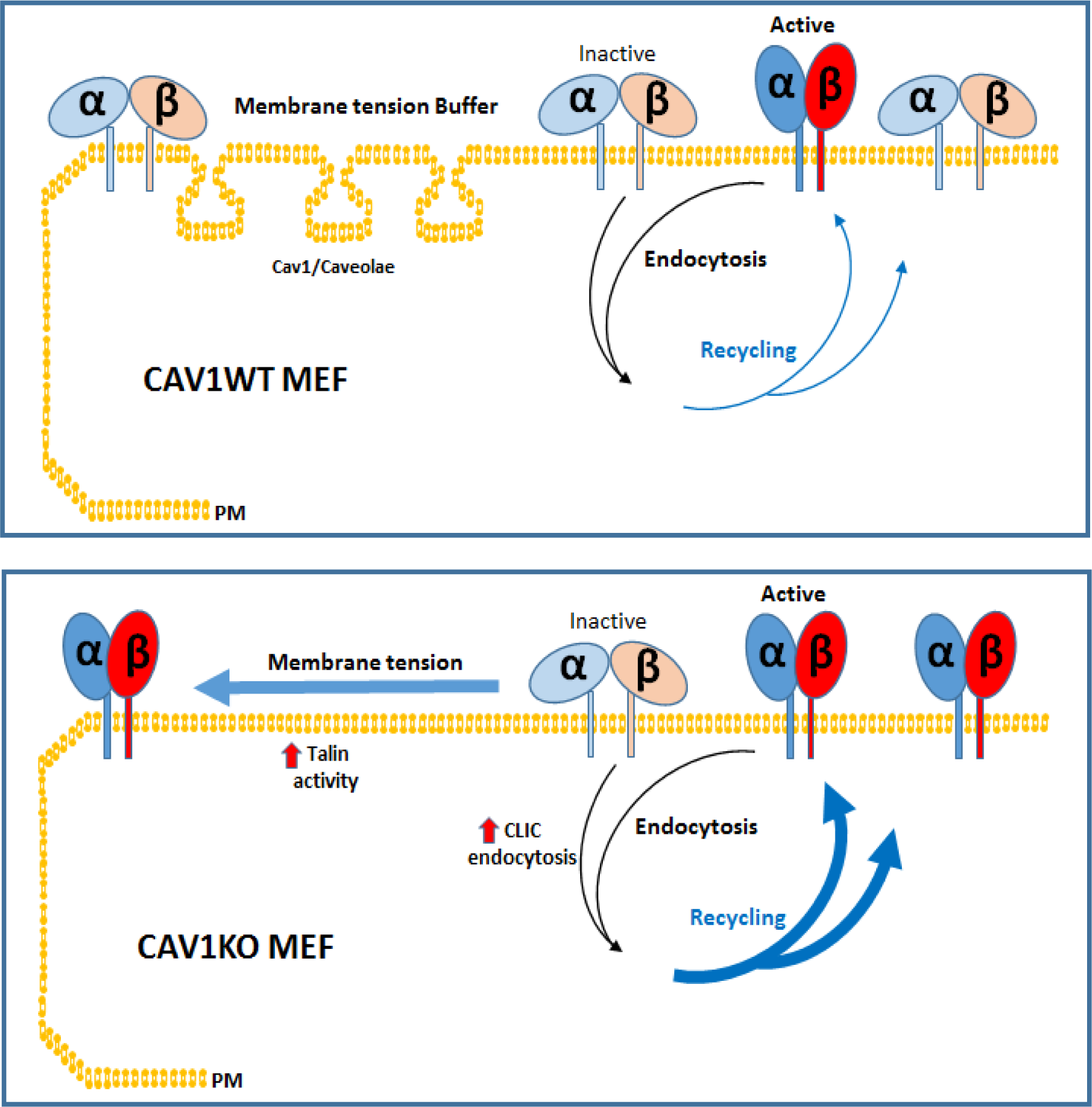
Caveolae adjust membrane tension to integrin mechanosensing by regulating integrin cycling and activation. Wild type MEFs adapt to membrane tension changes through the buffer system of caveolae, driving a physiological integrin mechanosensing (in this case α_5_β_1_-integrin). In the absence of caveolae, dysregulation of this response leads to increased PM tension, which accelerates integrin recycling and switches integrin from the inactive forms (light blue and light red integrin subunits) to the active forms (dark blue and dark red integrin subunits). Both increased β1-integrin recycling and activation is supported by increased talin activity in the absence of caveolae.

## Materials and methods

### Cloning, cells, and reagents

Caveolin-1 Flag was excised from pCDNA3.1 Cav1 with BamH1/EcoR1, klenow treated, and ligated into the klenow blunt-ended EcoR1 site of the GFP-expressing retroviral vector MIGR1. PTRF was excised from pIRES2-cavin1 EGFP with BglII/BamH1 and ligated into the BglII site of MIGR1. The C-terminally EGFP-tagged β1-integrin was developed by Prof. Martin Humphries (University of Manchester, UK) and described elsewhere^70^, and was requested through the Addgene public repository under ref. num. #69767.

All cells were cultured at 37°C and 5% CO_2_ in DMEM (Thermo Fisher Scientific™) supplemented with 10% fetal bovine serum (FBS) and 1% penicillin and streptomycin. Cav1KO MEFs were kindly provided by Michael Lisanti (Institute of Cancer Sciences, Manchester). All cell cultures were routinely checked for mycoplasma contamination. To deplete talin 2, cells were transfected with a plasmid encoding talin 2 shRNA and puromycin resistance^58^. Puromycin (2 µg/ml) was added to the cells 24 h after transfection and maintained for 4 days to select transfected cells. To deplete talin 1, talin 2 shRNA-stable cells were transfected with Tln1 esiRNA (EMU083531, Sigma Aldrich) or, for rescue experiments with the talin head, with Tln1 siRNA (Silencer® Select, Life technologies^TM^), which does not target the talin head domain. For rescue experiments, talin 2 shRNA-stable cells were co-transfected with Tln1 siRNA and EGFP-talin 1 head (Addgene plasmid no. 32856) or EGFP-talin 1 L325R (the mutant version, kindly provided by M. Ginsberg, UC San Diego, USA).

The following primary antibodies were used: rat monoclonal anti-mouse total β1-integrin (clone MB1.2, MAB1997 Millipore); rat monoclonal anti-mouse β1-integrin, activated (clone 9EG7, BD Pharmingen™); Alexa 488 conjugated anti-integrin β1, activated (clone HUTS-4, MAB2079-AF488 Millipore); rabbit polyclonal anti-mouse PTRF (Abcam); rabbit monoclonal anti-mouse caveolin-1 (Cell Signaling, #3238); mouse monoclonal anti-tubulin (Abcam, clone DM1A); rabbit monoclonal anti-CD147 (Invitrogen, clone JF1-045), mouse monoclonal anti-EEA-1 (BD transduction, clone 14) and mouse monoclonal anti-LBPA (Echelon Z-SLBPA). Secondary antibodies were Alexa Fluor®-488 goat anti-rat (Thermo Fisher Scientific™); Alexa Fluor®-647 goat anti-rat (Thermo Fisher Scientific™); Alexa Fluor®-488 phalloidin (Thermo Fisher Scientific™); HRP-linked anti-biotin from Cell Signaling (#7075); and Alexa Fluor®-647 phalloidin (Thermo Fisher Scientific™). EZ-Link™ SulfoNHS-SS-biotin was from Thermo Fisher Scientific (D21331), 2-mercaptoethanesulfonic acid (MESNA) and iodoacetamide from Sigma Aldrich (63707 and I1149), the Cdc42 inhibitor ML141 from Tocris Bioscience, and Alexa Fluor™ 647-Dextran from Thermo Fisher Scientific (D22914). Silencing of p190RhoGAP was as previously described^20^.

### PM fractionation and western blot analysis

MEFs were processed for PM isolation as described^71^. All steps were carried out at 4°C. Cells were first washed with cold-PBS 1X and pelleted by centrifugation at 14000xg for 5 minutes. Cells were then manually homogenized with 20 strokes of a PTFE head Tissue homogenizer (VWR®) and centrifuged at 1000xg for 10 minutes. The post-nuclear supernatant was collected and layered atop a 30% Percoll column. After centrifugation of the Percoll column at 84000xg for 30 minutes, the PM fraction was a visible band around 5.7 cm from the bottom of the centrifuge tube, was collected, further centrifuged at 105000g for 1 h to remove Percoll, separated by SDS-PAGE, and finally analyzed by western blot. Samples were immunoblotted with rabbit monoclonal anti-mouse caveolin-1 and rat monoclonal anti-mouse total β1-integrin (clone MB1.2, MAB1997 Millipore) as a loading control and a marker for PM fraction. Total cell lysates were separated by SDS-PAGE and analyzed by western blot with rabbit monoclonal anti-mouse Caveolin-1 and rabbit polyclonal anti-mouse PTRF (Abcam), with mouse monoclonal anti tubulin used as the loading control. Secondary antibodies were goat anti-mouse 800 and goat anti-rabbit 680. All membranes were scanned with the Odyssey imaging system (Li-COR®).

### Electron microscopy

MEFs were processed for electron microscopy using standard procedures. Briefly, cells were fixed for 1 hour with 2.5% glutaraldehyde in 100 mM cacodylate buffer, pH 7.4, and then post-fixed for 3 hours with 1% osmium tetroxide in 100mM cacodylate buffer, pH 7.4. The samples were dehydrated with acetone, embedded in Epon and sectioned. Ruthenium red (1 mg/ml) was added during fixing and post-fixing to stain the PM.

### Confocal and ground state depletion microscopy

Confocal images were obtained with an LSM 700 inverted confocal microscope (Carl Zeiss) fitted with a 63× 1.4 NA objective and driven by Zen software (Carl Zeiss). Superresolution imaging was performed with a GSD-TIRF microscope (Leica Microsystems). Samples were prepared according to standard procedures indicated by Leica Microsystems. The primary antibody was rabbit monoclonal anti-mouse caveolin-1 (1:100) and Alexa Fluor® 647 Fab1 fragment goat anti-rabbit (Jackson Immunoresearch; 1:100) was used as the secondary to further improve spatial resolution.

### Magnetic tweezers and reinforcement measurements

#### Bead coating

Carboxylated magnetic beads (Invitrogen) were mixed in a solution containing 500 µl 0.01 M sodium acetate pH 5, 0.75 mg Avidin (Invitrogen), and 4 mg EDAC (Sigma). Beads were incubated for 2h at RT and then washed in PBS and further incubated for 30 minutes in 1 ml 50 mM ethanolamine (Polysciences). The beads were then washed 3 times in PBS and left in PBS on a cold room rotator.

#### Force measurements

Magnetic tweezers experiments were performed as described^72, 73^. Briefly, carboxylated 3 μm magnetic beads (Invitrogen) were coated with biotinylated pentameric FN7-10 or ConA (Sigma Aldrich) mixed 1:1 with biotinylated BSA. For measurements, cells were first plated on coverslips coated with 10 μg/ml FN (Sigma) in Ringer’s solution (150 mM NaCl, 5 mM KCl, 1 mM CaCl2, 1 mM MgCl2, 20 mM HEPES, and 2 g/L glucose, pH 7.4) for 30 minutes. FN-coated beads were then deposited on the coverslips and allowed to attach to the cells. The tip of the magnetic tweezers device was then used to apply a force of 1 nN for 2 or 3 min to beads attached to cell lamellipodia. The apparatus used to apply force to the magnetic beads was as previously described^17^. The system was then mounted on a motorized 37 °C stage on a Nikon fluorescence microscope. DIC images and videos were recorded with a 60X objective linked to a CCD camera at a frequency of 250 frames/second.

### Magnetic twisting

Cells seeded on FN-coated coverslips were subjected to magnetic twisting as previously described^74^. After twisting, cells were fixed for 20 min in 4% paraformaldehyde (PFA) and then stained with 9EG7 antibody without permeabilization.

### Silica bead coating and staining

Carboxylated silica beads (3 μm, Kisker Biotech) were prepared as described above except that ConA-coated beads were incubated with biotinylated BSA previously labeled with an Alexa Fluor 555 protein labeling kit (Invitrogen). Unlabeled beads (FN-coated) and labeled beads (ConA-coated) were mixed in the same proportion (1:1). Cells were allowed to spread for 15 min and then fixed with 4% PFA, permeabilized with 0.1% Triton-X 100, and incubated at room temperature for 1 h with the indicated antibodies. Only beads at the cell periphery were analyzed (excluding cells in the ectoplasm-endoplasm border zone). To quantify fluorescence intensity, a 10-pixel-diameter ring was drawn around each selected bead using ImageJ. Mean fluorescence per area was normalized and plotted.

### Traction force microscopy

Traction force was measured as previously described^5^. Briefly, cells were seeded on polyacrylamide gels incorporating embedded fluorescent nanobeads. Cells were imaged by Phase contrast and embedded nanobeads by fluorescence. Cells were then trypsinized and bead position images were acquired in the relaxed gel state. Comparison of bead positions in gels with and without cells was used to obtain a gel deformation map^75, 76^. Images were obtained with a Nikon Eclipse Ti inverted microscope fitted with a 40x objective (Numerical Aperture = 0.6).

### Adhesion assay

Cell adhesiveness was assessed by seeding MEFs on 96-well plates coated with FN or ConA (both at 5 µg/ml) and incubating at 37°C for 30 minutes. Wells with no coating were included as negative controls. Cells were then fixed with methanol and stained with crystal violet (Sigma Aldrich). Wells were washed thoroughly to remove excess dye and were finally eluted with a mixture of 50% ethanol and 50% 0.1M sodium citrate (pH 4.2). The absorbance was read at 595 nm.

### FRAP

Cav1KO and wild type MEFs were transfected with the β1-GFP expression vector. Two pre-bleached events were acquired before bleaching by stimulation with the Nikkon scanner at 488 nm. Fluorescence recovery was monitored continuously until the intensity plateaued (approximately 1.5 minutes). Fluorescence during recovery was normalized to the pre-bleach intensity. Cells were cultured in DMEM (Thermo Fisher Scientific™) supplemented with 10% fetal bovine serum (FBS), and 1% penicillin and streptomycin.

### Endocytosis/recycling assay

β1-integrin kinetics were analyzed after biotin labeling of cell-surface integrins followed by a capture ELISA-based assay, using a modification of a previously described protocol^25^.

**Cell-surface integrin biotinylation** wild type or Cav1KO MEFs (5x10^5^) were seeded on 5 (endocytosis) or 4 (recycling) matrigel-coated plates* (labeled Total, 0 minutes, 2 minutes, 5 minutes, and 10 minutes for endocytosis and 0 minutes, 1 minute, 3 minutes and 5 minutes for recycling). The cells were incubated in complete DMEM (Thermo Fisher Scientific) for 2 hours at 37°C, which was the shortest spreading time for cells to stand the assay conditions, and also matches the magnetic tweezers measurement experimental time frame. The plates were then placed on ice, washed twice with ice-cold PBS and incubated for 40 minutes at 4°C with 0.25 mg/ml of EZ-Link™ SulfoNHS-SS-Biotin in Hank’s balanced salt solution (Sigma Aldrich). After two further washes with ice-cold PBS, the plates were labeled for the appropriate time points and processed as described below. *Matrigel was used instead of FN to mimic a more physiological environment. While different coatings can clearly affect integrin trafficking, matrigel contains, among other extracellular components, collagen and certain levels of fibronectin, which we have shown to increase Cav1KO adhesion as compared to wild type MEFs (see Suppl. Figure 1B and 1C). Importantly, cells were allowed to spread for 2 hours before starting the assay in complete medium, thus ensuring an additional supply of FN.

#### Endocytosis

The 2, 5, and 10 minute plates were incubated at 37°C with 2 ml of pre-warmed DMEM (without FBS) for the indicated times. The Total and 0 minute plates were placed on ice with 2 ml ice-cold DMEM (without FBS). All plates except Total were then washed twice with ice-cold PBS and incubated for 40 minutes at 4°C with MESNA-containing buffer (Sigma Aldrich) to remove remaining surface-associated biotin. All the plates were then washed twice with ice-cold PBS and incubated with iodoacetamide for 10 minutes at 4°C. After washing again with ice-cold PBS, cells were lysed and processed for ELISA.

#### Recycling

The 0, 1, 3, and 5 minute plates were incubated at 37°C with 2 ml of pre-warmed DMEM (without FBS) for 10 minutes (to allow time for endocytosis). The plates were then washed twice with ice-cold PBS and incubated for 40 minutes at 4°C with MESNA-containing buffer to remove remaining surface-associated biotin. The 1, 3, and 5 minute plates were incubated at 37°C with 2 ml of pre-warmed DMEM (without FBS) for the indicated times; the 0 minute plate was placed on ice with 2 ml ice-cold DMEM (without FBS). At the end of the incubation, all plates were washed twice with ice-cold PBS and incubated again with MESNA-containing buffer to remove biotin-labeled integrins that had recycled to the PM. Plates were finally washed with ice-cold PBS, incubated with iodoacetamide for 10 minutes at 4°C, lysed, and processed for ELISA.

#### ELISA-based assay

96-well ELISA plates were coated overnight at 4°C with anti-mouse total β1-integrin (Millipore) or anti-mouse β1-integrin, activated (BD Pharmingen™). The plates were then washed three times with solution A (0.02% Tween-20 in PBS), blocked for 1 hour at room temperature with solution B (0.02% Tween-20, 1% BSA in PBS), and incubated with cell lysates from endocytosis or recycling assays for 2 hours at room temperature or overnight at 4°C. After washing three times with solution A, the plates were incubated for 1 hour at room temperature with anti-biotin HRP-linked antibody. Plates were then washed three more times with solution A, and integrin was detected by TMB reaction (Sigma Aldrich). Endocytosis results were normalized by dividing with the signal from Total wells; graphs represent the progressive increase in the amount of total or active β1-integrin. Recycling results were normalized by dividing with the signal from 0-minute wells; graphs represent the β1-integrin remaining inside cells, so that negative curves indicate an increase in the recycling rate.

**To determine total cell-surface integrin**, cells were processed as in the total plates. For Rab DNs and Tln siRNA experiments, cells were plated 48h before the experiment avoiding re-plating to prevent losing lesser adherent cells. For cholesterol loading experiments, cells were treated with U18666A 2ugr/ml or LDL 100ugr/ml for 24h before surface integrin quantification.

### Hypoosmotic treatment

Cells were cultured for the indicated time points at 37°C in DMEM diluted as indicated with distilled water. The cells were then immediately fixed for 15 minutes by adding an equal volume of 8% PFA (yielding a final PFA concentration of 4%) and then stained with 9EG7 antibody without permeabilization

### Endocytosis experiments

For dextran endocytosis: cells were first incubated for 1 hour at 4°C (to prevent endocytosis) with β1-Alexa 488 conjugated antibody, activated. Cells were then washed twice with PBS and incubated for 3 min at 37°C with 1 mg/ml Alexa Fluor™ 647-Dextran (Invitrogen, REF D22914) without preincubation or acid stripping. Cells were then fixed with 4% PFA and analyzed by confocal microscopy. For other endocytosis experiments please see extended material and methods. Colocalization was analyzed using the plugin Coloc 2 (Fiji^77^).

### Extracellular staining

To analyze cell-surface β1-integrin, cells were fixed for 20 min with 4% PFA and then stained with 9EG7 antibody without permeabilization.

### Statistical analysis

Data are presented as mean ± s.e.m. unless otherwise indicated. Mean values were compared by two-tailed paired Student *t* test unless otherwise indicated. Differences were considered statistically significant at P < 0.05 (*), < 0.01 (**), and < 0.001 (***).

## Acknowledgements

We thank Dr. Miguel Sánchez for critical reading of the manuscript and Simon Bartlett for scientific editing. We also thank Verónica Labrador Cantarero and Antonio M. Santos Beneit from Microscopy Unit (CNIC) for macro development and video editing and Dr. Martin Humphries (The University of Manchester), Dr. Michael Lisanti (Institute of Cancer Sciences, Manchester), Dr. M. Ginsberg (UC San Diego, USA) and Dr. Cristina Clemente Toribio for kindly providing reagents and cells.

This project received funding from the European Union Horizon 2020 Research and Innovation Programme through Marie Sklodowska-Curie grant 641639; and grants from the Spanish Ministry of Economy, Industry and Competitiveness (MINECO; SAF2011-25047, SAF2014-51876-R, SAF2017-83130-R, IGP-SO grant MINSEV1512-07-2016, CSD2009-0016, BFU2016-81912-REDC), the Worldwide Cancer Research Foundation (#15 -0404), and the Asociación Española Contra el Cáncer foundation (PROYE20089DELP) all to MAdP. MAdP is member of the Tec4Bio consortium (ref. S2018/NMT¬4443; ‘Actividades de I+D entre Grupos de Investigación en Tecnologíaś, Comunidad Autónoma de Madrid/FEDER, Spain), co-recipient with PR-C of grants from Fundació La Marató de TV3 (674/C/2013 and 201936-30-31), and coordinator of a Health Research consortium grant from Fundación Obra Social La Caixa (AtheroConvergence, HR20-00075). The CNIC Unit of Microscopy and Dynamic Imaging is supported by FEDER “Una manera de hacer Europa” (ReDIB ICTS infrastructure TRIMA@CNIC, Ministerio de Ciencia e Innovación (MCIN)). The CNIC is supported by the Instituto de Salud Carlos III (ISCIII), the Ministerio de Ciencia, Innovación y Universidades (MCNU) and the Pro CNIC Foundation, and is a Severo Ochoa Center of Excellence (SEV-2015-0505)

## Author contributions

Conceived and designed the experiments: F.N.L., P.R.C. and M.A.dP. Performed the experiments: F.N.L., D.M.P., A.G.G., A.E.A., V.I.S. and S.S.P. Analyzed the data: F.N.L. and D.M.P. Contributed to manuscript drafting: F.N.L., X.T, P.R.C. and M.A.dP.

## Supplementary Information

Figure legends, table and videos

**Supplementary Figure 1. Related to Figure 3.**
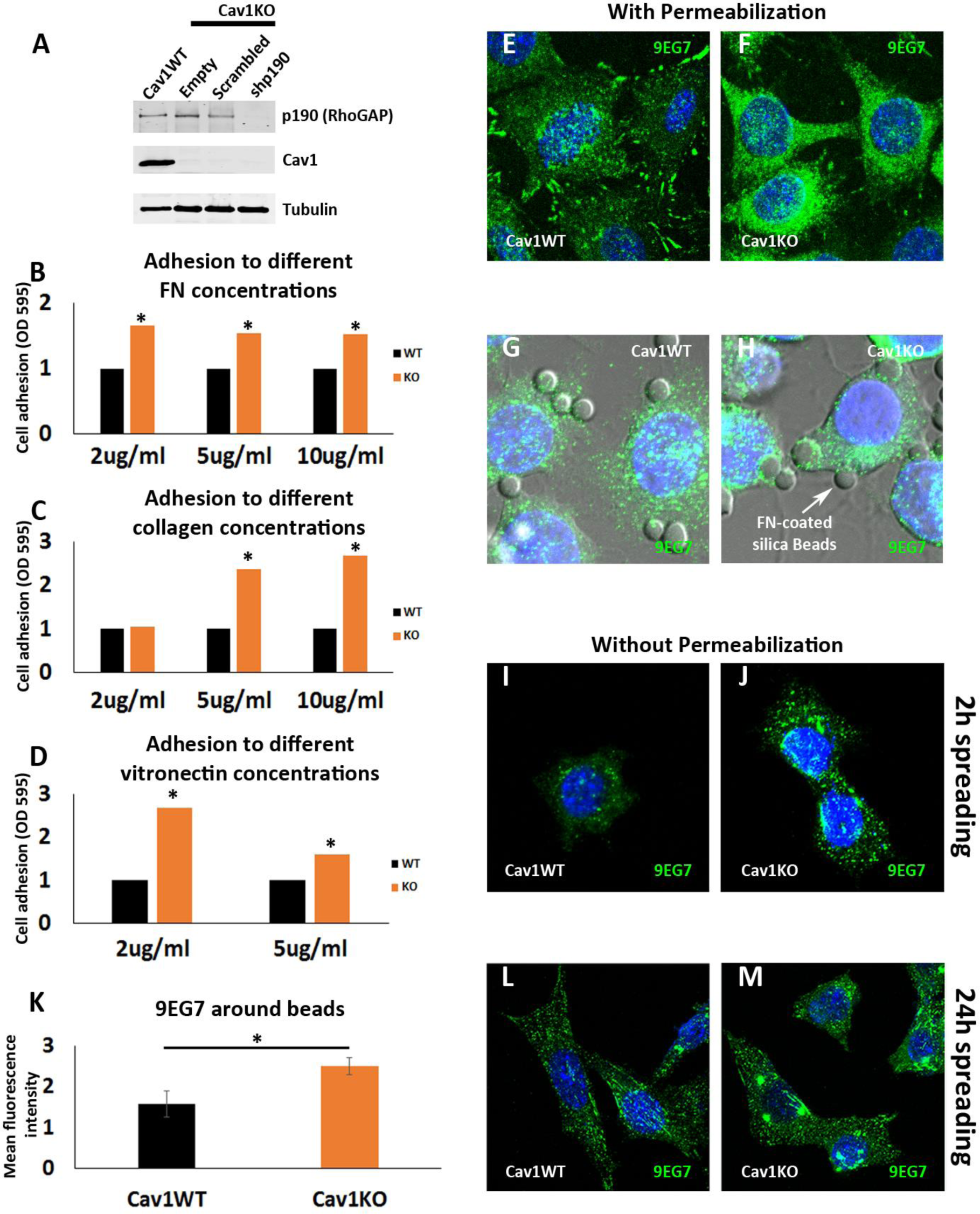
(A) Western blot of total lysates from wild type cells, and Cav1KO cells reconstituted with either empty vector, scrambled shRNA or shRNA against p190RhoGAP. Samples were immunoblotted for p190RhoGAP, Cav1, and tubulin (loading control, 50kDa). (B, C and D) Relative adhesion of wild type and Cav1KO MEFs to different concentrations of fibronectin (B), collagen (C) and vitronectin (D). Measurements (absorbance, Optical density, OD, at 595nm from retained crystal violet dye, see material and methods) were normalized to values from wild type counterparts. (E and F) Confocal microscopy images of active β1-integrin immunostaining (9EG7 antibody, green) in permeabilized wild type (E) MEFs or Cav1KO (F) MEFs. DAPI counterstain is shown in blue. (G and H) Confocal microscopy images of active β1-integrin immunostaining (9EG7 antibody, green) in wild type (G) or Cav1KO (H) MEFs incubated with FN-coated silica beads (arrow). DAPI counterstain is shown in blue. (I, J, L and M) Confocal microscopy images of active β1-integrin immunostaining (9EG7 antibody, green) in non-permeabilized wild type MEFs (I and L) and Cav1KO MEFs (J and M) after the indicated spreading times. DAPI is shown in blue. (K) Normalized mean bead-bound 9EG7 fluorescence intensity in wild type and Cav1KO MEFs; n ≥ 40 beads per genotype. Statistical comparisons were by *t*-test, with significant between-group differences denoted \**P*<0.05. Figure 3**-figure supplement 1-source data 1 and 2** includes the full raw unedited blot (1 corresponding to Suppl. Figure 1A) and the uncropped blot with the relevant bands labelled (2 corresponding to Suppl. Figure 1A). **See also Raw Data Suppl. Figures 1 and 2** which includes the raw data of experiments from Suppl. Figure 1B, 1C, 1D and 1K.

**Supplementary Figure 2. Related to Figure 4.**
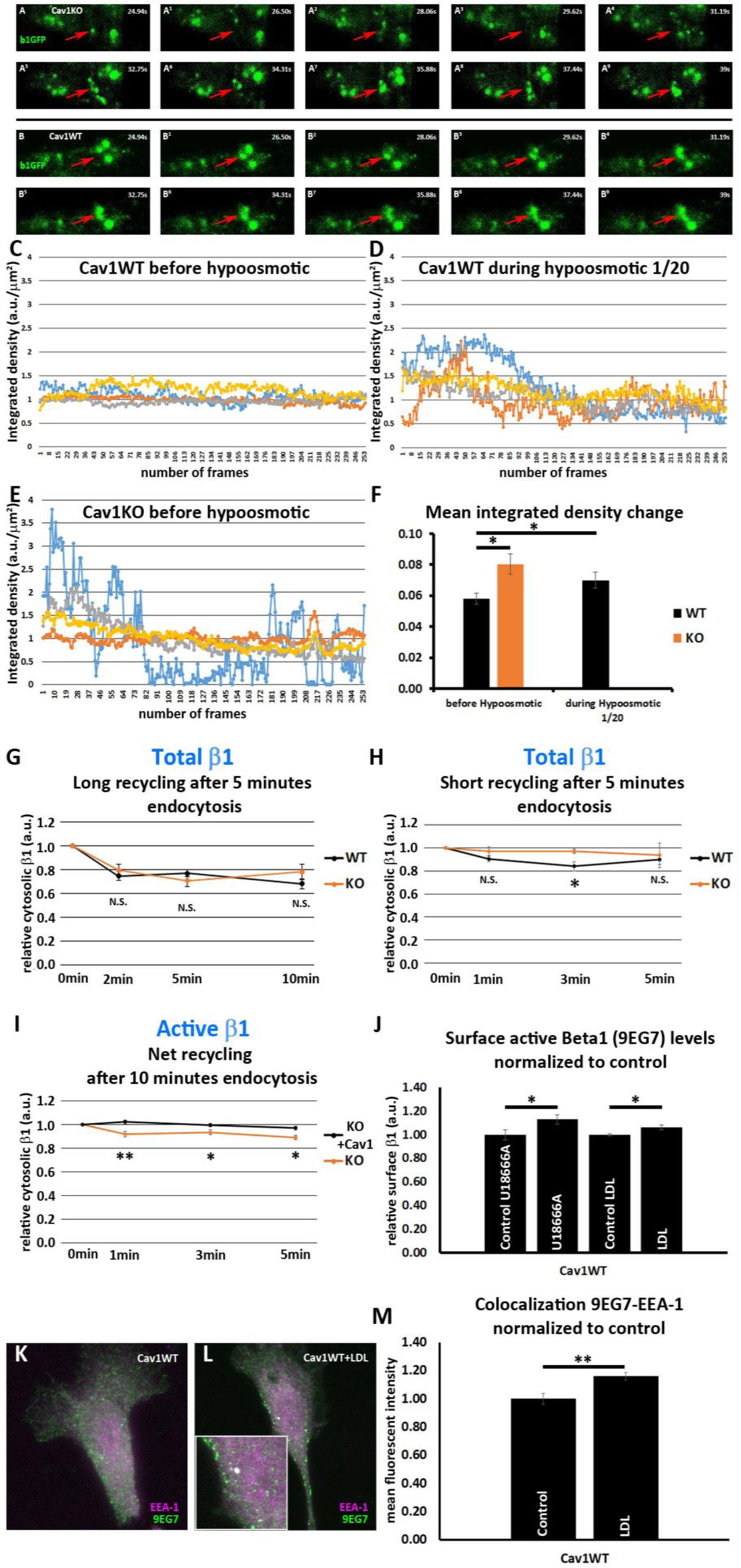
(A-A^9^) TIRFm image sequence (∼1.5-second intervals) of Cav1KO MEFs transfected with β1-integrin-GFP (green). Arrows indicate the dynamic behavior of GFP-positive vesicles in the TIRF plane. (B-B^9^) TIRFm image sequence (∼1.5-second intervals) of wild type MEFs transfected with β1-integrin-GFP (green). Arrows indicate the stability of GFP-positive vesicles in the TIRF plane. (C-F) Quantification of TIRFm videos, represented as the change in normalized fluorescence integrated density, a measure of the mean fluorescence intensity in each video frame (see material and methods for details). (G and H) Net recycling after 5 minutes endocytosis of total β1-integrin in wild type and Cav1KO MEFs over (G) long time-point set or (H) short time-point set (net recycling is expressed as internal biotinylated β1 integrin at each time point normalized to time point 0 (which contains all the biotinylated β1 integrin internalized after 5 minutes of endocytosis, see Materials and Methods); n ≥ 6 recycling assays per genotype. (I) Net recycling after 10 minutes endocytosis of active β1-integrin in Cav1KO and Cav1KO reconstituted with Cav1 MEFs at the time points indicated. Net recycling is expressed as internal biotinylated β1 integrin at each time point normalized to time point 0 (which contains all the biotinylated β1 integrin internalized after 10 minutes of endocytosis, see Materials and Methods); n = 6 recycling assays per genotype. (J) Quantification of the impact on surface active β1-integrin levels in wild type MEFs of treatment with U18666A 2ugr/ml or LDL 100ugr/ml for 24h. Values are expressed as normalized to control conditions across experiments; n ≥ 18 replicates from at least 6 independent experiments. (K and L) Confocal microscopy images of wild type MEFs, stained for active β1-integrin (9EG7 antibody, green) and EEA1 (magenta), either non-treated (K) or treated with 100ug/ml LDL for 24h (L). (M) Quantification of colocalization between EEA1 and 9EG7, expressed as Pearson’s correlation coefficient; n ≥ 25 cells per condition. Statistical significance of differences across indicated conditions was assessed by *t*-test: * *p*<0.05 P\*\**p*<0.01; N. S., non-significant. **See also Raw Data Suppl. Figures 1 and 2** which includes the raw data of experiments from Suppl. Figure 2F, 2G-2J and 2M.

**Supplementary Figure 3. Related to Figure 5.**
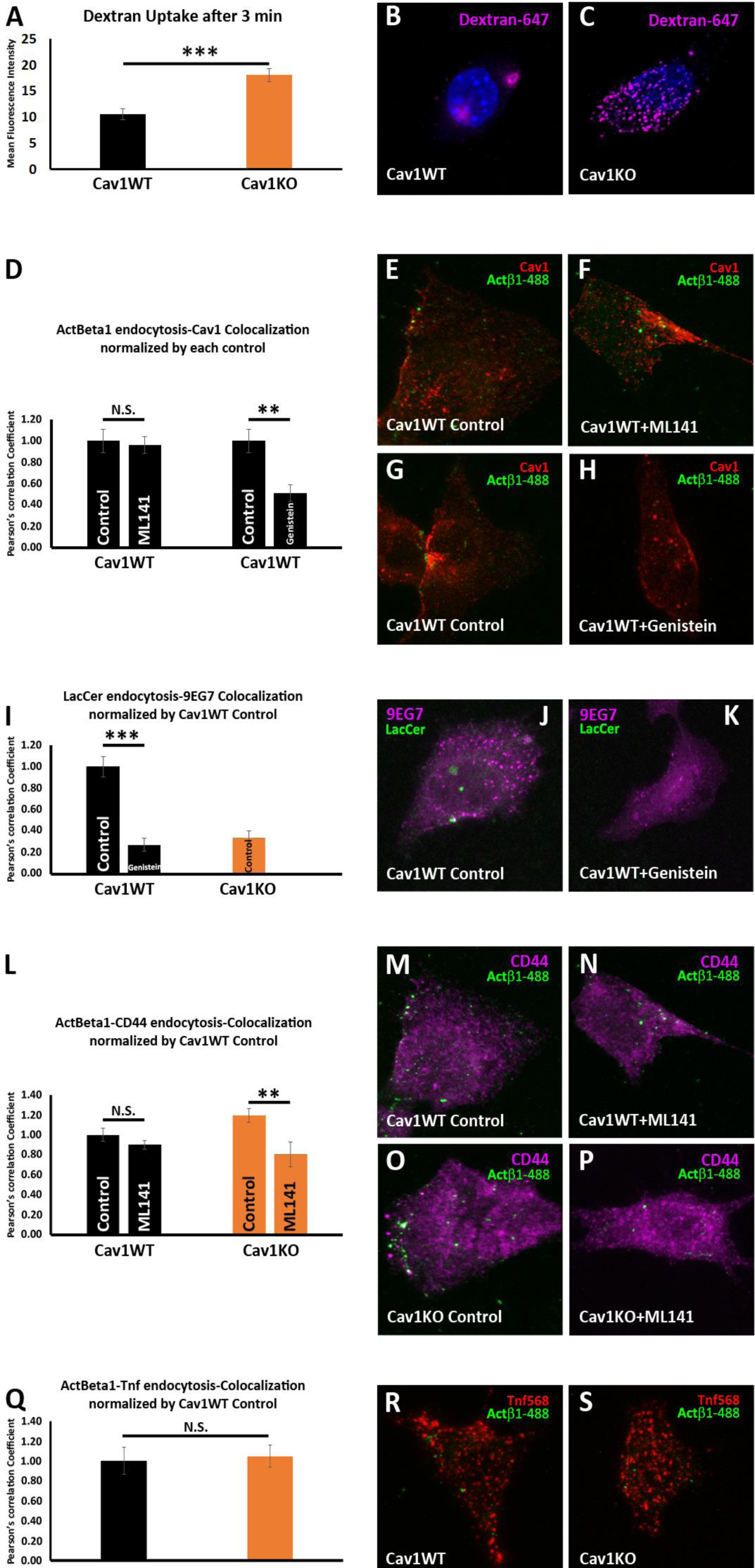
(A) Quantification of Dextran-647 mean fluorescent intensity after 3 minutes of endocytosis in wild type and Cav1KO MEFs; n ≥ 25 cells per genotype. (B and C) Representative confocal microscopy images of (B) wild type MEFs and (C) Cav1KO MEFs after 3 minutes of Dextran-647 endocytosis. DAPI counterstain in shown in blue. (D) Quantification of colocalization between active β1-Alexa 488 and Caveolin1 in wild type MEFs non-treated (control) or treated with ML141 inhibitor 10uM for 30 min or genistein 200uM for 2h, expressed as Pearson’s correlation coefficient normalized by each control; n ≥ 10 cells per genotype. (E-H) Representative confocal microscopy images of wild type MEFs either non-treated (E and G, control) or treated with ML141 10uM for 30 min or genistein 200uM for 2h before incubation with anti-active β1-Alexa 488 antibody (green) for 1 hour at 4°C followed by 3 minutes endocytosis at 37°C. Caveolin1 staining is shown in red (Cav1). (I) Quantification of colocalization between BODIPY-LacCer and active β1-integrin (9EG7) in wild type MEFs either non-treated (control) or treated with genistein 200uM for 2h and Cav1KO MEFs non-treated, expressed as Pearson’s correlation coefficient normalized by wild type control; n = 15 cells per genotype. (J and K) Representative confocal microscopy images of wild type MEFs either non-treated (J, control) or treated with genistein 200uM for 2h before incubation with BODIPY-LacCer 5uM (green) for 1 hour at 4°C followed by 3 minutes endocytosis at 37°C. BODIPY-LacCer remaining at the PM was then removed by back exchange at 4°C (see material and methods for more details). Active β1-integrin staining is shown in magenta (9EG7). (L) Quantification of colocalization between active β1-Alexa 488 and CD44 in wild type and Cav1KO MEFs either non-treated (control) or treated with ML141 inhibitor, expressed as Pearson’s correlation coefficient normalized by wild type control; n ≥ 30 cells per genotype. (M-P) Representative confocal microscopy images of wild type and Cav1KO MEFs either non-treated (M and O, control) or treated with ML141 inhibitor 10uM for 30 minutes before incubation with anti-active β1-Alexa 488 antibody (green) and anti-CD44 antibody (magenta) for 1 hour at 4°C followed by 3 minutes endocytosis at 37°C. (Q) Quantification of colocalization between active β1-Alexa 488 and Transferrin-568 (Tnf) in wild type and Cav1KO MEFs, expressed as Pearson’s correlation coefficient normalized by wild type control; n = 20 cells per genotype. (R and S) Representative confocal microscopy images of wild type and Cav1KO MEFs incubated with anti-active β1-Alexa 488 antibody (green) and Tnf-568 (magenta) for 1 hour at 4°C followed by 3 minutes endocytosis at 37°C. Statistical significance of differences across indicated conditions was assessed by *t*-test: * *p*<0.05 P\*\**p*<0.01; \*\*\**p*<0.001 N. S., non-significant.**See also Raw Data Suppl. Figures 3 and 4** which includes the raw data of experiments from Suppl. Figure 3A, 3D, 3I, 3L and 1Q.

**Supplementary Figure 4. Related to Figure 6.**
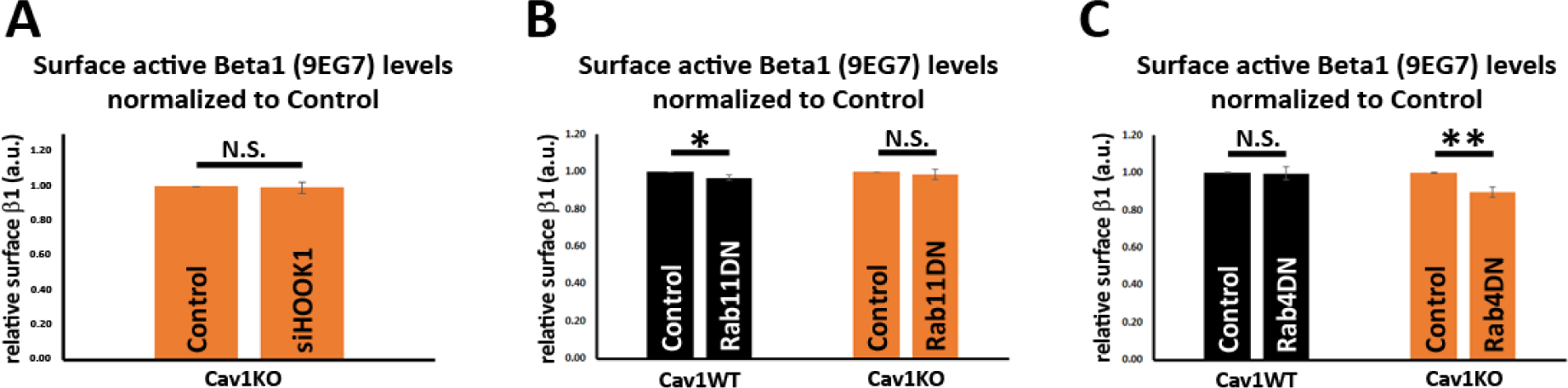
(A-C) Quantification of the impact on surface active β1-integrin levels in wild type or Cav1KO cells of: siRNA-mediated depletion against HOOK1 for 72h (A); expression of Rab4 S22N dominant negative mutant for 48h (B); expression of Rab11 N124I dominant negative mutant for 48h (C). Values are expressed as normalized to control conditions across experiments; n ≥ 5 independent experiments. Statistical significance of differences across indicated conditions was assessed by *t*-test: * *p*<0.05 P\*\**p*<0.01; \*\*\**p*<0.001 N. S., non-significant. **See also Raw Data Suppl.** Figures 3 and 4 which includes the raw data of experiments from Suppl. Figure 4A-4C.

**Supplementary Figure 5. Related to Figure 7.**
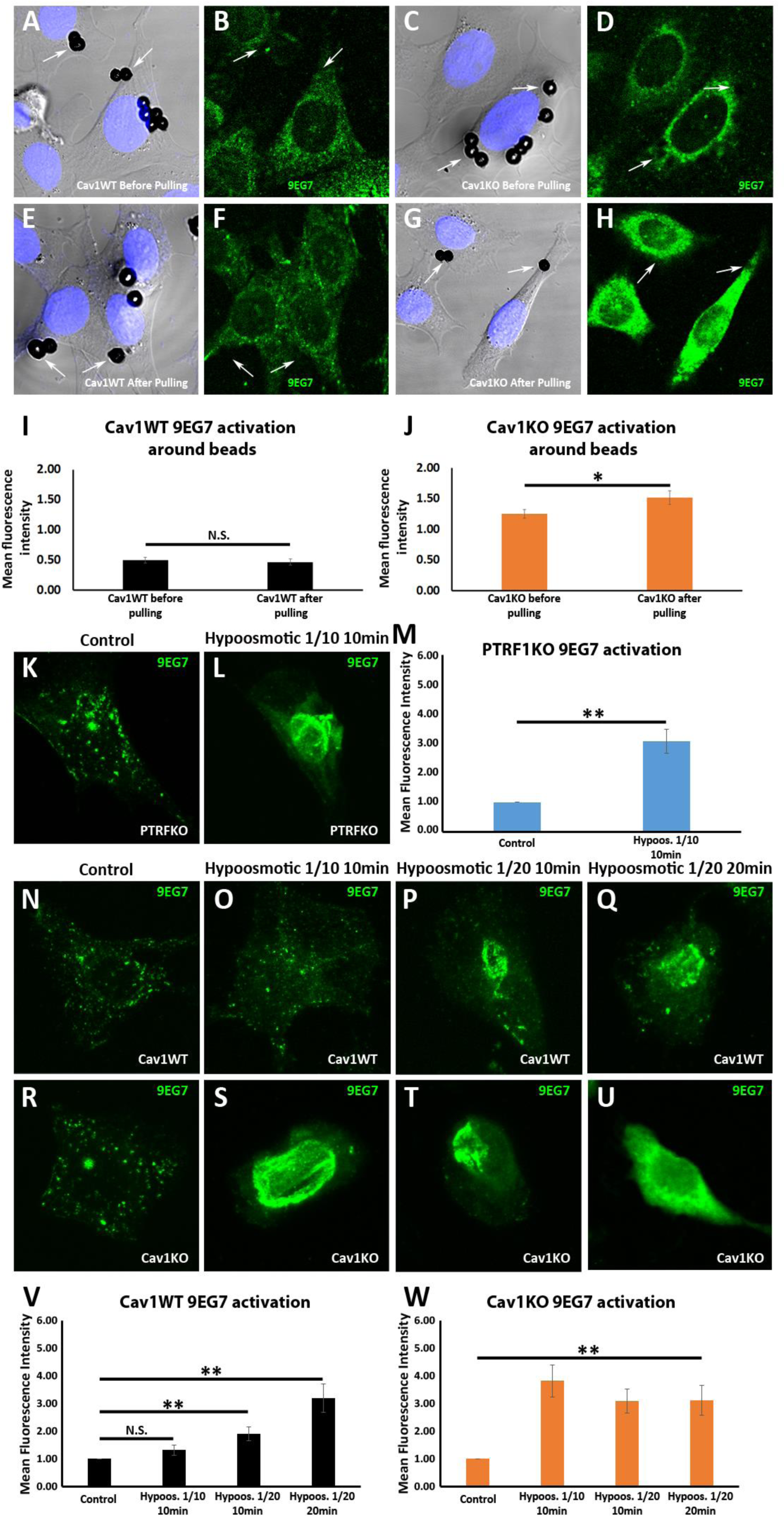
(A-H) Confocal microscopy images of wild type and Cav1KO MEFs before (A and B, C and D) and after (E and F, G and H) magnetic puling with FN-coated ferromagnetic beads. Active β1-integrin is shown in green (9EG7 staining) and DAPI in blue. Arrows indicate staining around beads. (I and J) Mean bead-bound 9EG7 fluorescence intensity normalized by bead area of (I) wild type MEFs and (J) Cav1KO MEFs before and after magnetic pulling; n ≥ 56 beads per genotype and condition. (K and L) Active β1-integrin immunostaining (9EG7, green) in PTRFKO MEFs cultured in standard (control) or hypoosmotic medium. (M) 9EG7 mean fluorescence intensity in control and hypoosmotic shock-exposed PTRFKO MEFs; n ≥ 30 cells per condition. (N-U) Active β1-integrin immunostaining (9EG7, green) in wild type (N-Q) or Cav1KO MEFs (R-U) cultured in standard (control) or in the different hypoosmotic conditions and time-points indicated; n ≥ 28 cells per genotype and condition. (V and W) 9EG7 mean fluorescence intensity in control and hypoosmotic shock-exposed wild type (V) or Cav1KO (W) MEFs at the hypoosmotic and time-points indicated. All immunostainings in this figure were performed following the extracellular staining (see material and methods for more details). Statistical comparisons were by *t*-test, with significance between groups denoted \**P*<0.05; \*\**p*<0.01; N. S., non-significant. **See also Raw Data Suppl. Figures 5 and 6** which includes the raw data of experiments from Suppl. Figure 5I, 5J, 5M, 5V and 5W.

**Supplementary Figure 6. Related to Figure 7.**
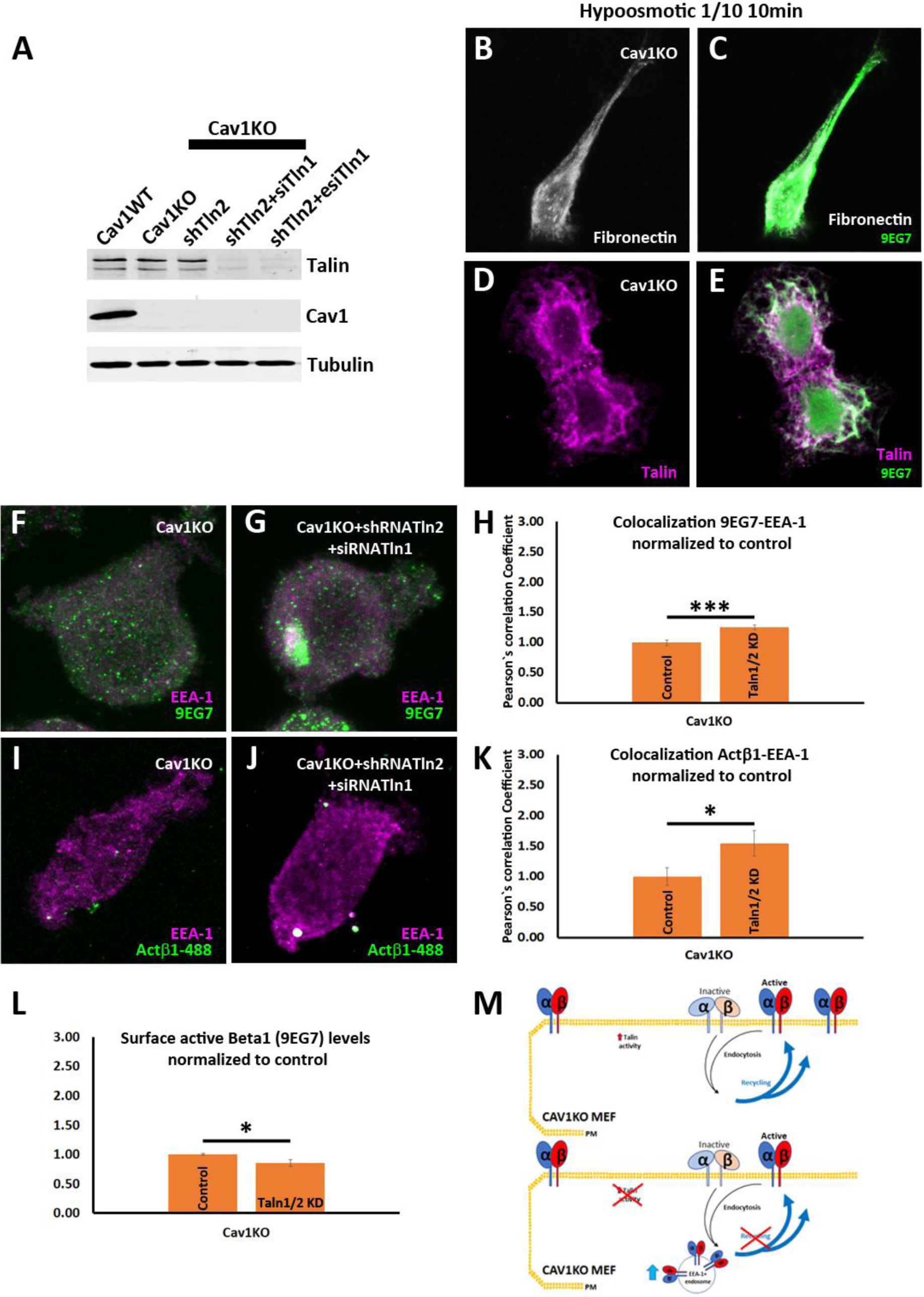
(A) Western blot analysis of total lysates from wild type and Cav1KO MEFs, either non-treated (Cav1KO), or treated with either shRNA against Talin2 alone (shTln2) or a combination of shRNA against Talin2 and siRNA against Talin1 (shTln2+siTln1 or shTln2+esiTln1, see material and methods for details). Membranes were immunoblotted for Talin, Cav1, and tubulin (loading control, 50kDa). (B-E) Confocal microscopy images of Cav1KO MEFs culture in hypoosmotic medium (diluted 1:10) for 10 minutes at 37°C, stained for active β1 integrin (9EG7 antibody, green) and Fibronectin-FITC (shown in grey, B and C) or Talin (shown in magenta, D and E). (F and G) Confocal microscopy images of Cav1KO MEFs, either non-treated (F), or treated with a combination of shRNA against Talin2 and siRNA against Talin1 (G), stained for active β1 integrin (9EG7 antibody, green) and EEA-1 (shown in magenta). (H) Quantification of colocalization between 9EG7 and EEA-1 normalized to control, expressed as Pearson’s correlation coefficient; n = 15 cells per condition. (I and J) Confocal microscopy images of Cav1KO MEFs, either non-treated (I), or treated with a combination of shRNA against Talin2 and siRNA against Talin1 (J) for 48h before incubation with anti-active β1-Alexa 488 antibody (green) for 1 hour at 4°C followed by 3 minutes endocytosis at 37°C. Cells were stained for EEA-1 (shown in magenta). (K) Quantification of colocalization between active β1-Alexa 488 and EEA-1 normalized to control, expressed as Pearson’s correlation coefficient; n ≥ 14 cells per condition. (L) Quantification of the impact on surface active β1-integrin levels in Cav1KO cells of a combination of shRNA against Talin2 and siRNA against Talin1. Values are expressed as normalized to control; n = 15 replicates from at least 5 independent experiments. Statistical comparisons were by *t*-test, with significance between groups denoted \**P*<0.05; \*\*\**p*<0.001. (M) Schematic representation showing that talin activity not only regulates β1 integrin activation but also supports increased recycling in Cav1KO MEFs. **Figure 7-figure supplement 6-source data 1 and 2** includes the full raw unedited blot (1 corresponding to Suppl. Figure 6A) and the uncropped blot with the relevant bands labelled (2 corresponding to Suppl. Figure 6A). **See also Raw Data Suppl.** Figures 5 and 6 which includes the raw data of experiments from Suppl. Figure 6H, 6K, 6L.

## Supplementary table 1. Related to Figure 2

Traces of beads movement shown in Figure 2C. They represent two examples of the data used to generate the reinforcement increment shown in Figure 2G.

## Supplementary Video 1. Related to Figure 2

Example magnetic tweezers experiment. Application of a pulsed magnetic force (1Hz, 1nN) causes the tip of the magnetic tweezers (black shadow to the right) to pull a magnetic bead attached to the cell surface (observed by DIC microscopy).

## Supplementary Video 2. Related to Figure 2

Molecular representation video of the type of forces generated by concanavalin A-coated beads. Force is exerted on the plasma membrane.

## Supplementary Video 3. Related to Figure 2

Molecular representation video of the type of forces generated by FN-coated beads. Force is exerted on integrins.

## Supplementary Video 4. Related to Figure 4

TIRFm video of a Cav1WT MEF transfected with β1-gfp expression vector.

## Supplementary Video 5. Related to Figure 4

TIRFm video of a Cav1KO MEF transfected with β1-gfp expression vector.

## Supplementary Video 6. Related to Figure 4

TIRFm video of a Cav1WT MEF transfected with β1-gfp expression vector, before hypoosmotic treatment.

## Supplementary Video 7. Related to Figure 4

TIRFm video of a Cav1WT MEF transfected with β1-gfp expression vector, during hypoosmotic treatment (1:20 DMEM dilution).

## Supplementary Video 8. Related to Figure 7

Example magnetic twisting experiment. Note how beads move under the oscillating magnetic field.

## Extended Material and methods

### Reagents

The following primary antibodies were used: mouse monoclonal anti-p190RhoGAP (Upstate, clone D2D6, 1:1000); rabbit monoclonal anti-mouse Caveolin-1 (Cell signaling®, 1:1000); mouse monoclonal anti-alpha tubulin (ab7291, Abcam, 1:10.000), mouse monoclonal anti-Talin (Sigma Aldrich, clone 8d4, 1:200) and mouse anti-CD44 (clone 5035-41.1D, Novus Biologicals) Fibronectin and LDLs were obtained by purification from blood donors and were conjugated with FITC (Thermo Fisher) or used directly, respectively. U18666A was from sigma (U3633).

### Total internal reflection fluorescent microscopy videos

TIRF microscopy was performed with a Leica AM TIRF MC microscope. TIRFm movies were acquired with a 100X_1.46 VNA oil-immersion objective at 488 nm excitation and an evanescent field with a nominal penetration depth of 100 nm. Images were collected with an ANDOR iXon CCD at 300 ms per frame. Quantification of TIRF videos show normalized fluorescence integrated density (IntDen) over frames (Suppl. Figure 2C-2E). Graph represents the mean of the difference between normalized fluorescence integrated density of adjacent frames (framex-framex-1) (Suppl. Figure 2F). IntDen was calculated as in the following formula: IntDen= Raw Integrated density (sum of pixel values in selection) x (Area in scaled units)/ (Area in pixels)), which was then normalized by the IntDen mean of all frames analyzed.

### Endocytosis experiments

**For caveolar uptake,** cells were treated, when indicated, with genistein 200uM for 2h before incubation with BODIPY-LacCer 5uM for 1 hour at 4°C followed by 3 minutes endocytosis at 37°C. BODIPY-LacCer (Invitrogen, REF B34402) remaining at the PM was then removed by back exchange at 4°C following a previous protocol^1^. Cells were then fixed with 4% PFA, stained and analyzed by confocal microscopy.

**For clathrin-independent endocytosis** (CLIC), cells were treated, when indicated, with ML141 inhibitor 10uM (Tocris, REF 4266) for 30 min before incubation with anti-active β1-Alexa 488 antibody and anti-CD44 antibody for 1 hour at 4°C followed by 3 minutes endocytosis at 37°C. Cells were then acid stripped to remove surface staining, fixed with 4% PFA, stained and analyzed by confocal microscopy.

**For clathrin endocytosis**, cells were incubated with anti-active β1-Alexa 488 antibody and Tnf-568 (Invitrogen, REF T23365) for 1 hour at 4°C followed by 3 minutes endocytosis at 37°C. Cells were then acid stripped to remove surface staining, fixed with 4% PFA, stained and analyzed by confocal microscopy.. Colocalization was analyzed using the plugin Coloc 2 (Fiji^2^).

